# Two phytohormones synergistically induce parasitic weeds seed germination via KAI2d receptors

**DOI:** 10.64898/2025.12.31.697154

**Authors:** Taiki Suzuki, Tomoya Ishikawa, Marco Bürger, Masahiko Otani, Koji Miyamoto, Wendi Jiang, Hanae Kaku, Naoki Kitaoka, Hideyuki Matsuura, Michio Kuruma, Kotaro Nishiyama, Yoshiya Seto

## Abstract

Root parasitic plants can severely decrease global crop production. The germination of their seeds is induced by host root-derived strigolactones (SLs). Considering this unique germination system, a “suicidal germination” method has been proposed to control root parasitic plants. However, this method requires the cost-effective production of germination inducers. In this study, we determined that jasmonates and SLs can synergistically induce the seed germination of root parasitic plants (*Orobanche minor* and *Striga hermonthica*). Biochemical analyses indicated that jasmonates and SLs cooperatively activate multiple divergent KARRIKIN INSENSITIVE 2 receptors. Our findings have elucidated the host recognition systems associated with these highly duplicated promiscuous receptors, with potential implications for developing a new strategy to protect crops against root parasitic plants.

## Main Text

Root parasitic plants obtain water and nutrients from host plants via their xylem tissue, which is connected to the host vasculature. Parasitic plants in the family Orobanchaceae, such as *Striga* and *Orobanche* spp., cause significant crop losses worldwide and threaten food security (*1*). Their seeds remain dormant for decades in soil, but they can germinate if exposed to host root-derived small molecules, namely strigolactones (SLs), which are biosynthesized as plant hormones and rhizosphere signaling molecules for arbuscular mycorrhizal symbiosis (*2–5*). Although unique germination system is essential for initiating their lifecycle near the indispensable host plants, it leads to the development of an effective strategy, which is called “suicidal germination”, to control root parasitic plants. Thus, root parasitic plants may be eliminated from infested fields by artificially inducing the germination of their seeds in the absence of host plants using germination-inducing compounds, including SL-like molecules. However, the feasibility of this approach is restricted by the difficulty of producing germination inducers in sufficient quantities.

A fermentation-based method for the quantitative production of the plant hormone gibberellin (GA) was developed using *Gibberella fujikuroi*, a GA-producing phytopathogen (*6*). Recently, *Exiguobacterium* R2567 was revealed to produce cyclo(Leu-Pro), which is a compound with SL-like activity that can regulate the number of tillers in rice plants (*7*). Considering there are many reports describing bacteria and fungi that can produce plant hormones or their mimics, we speculated that microorganism metabolites may include SLs or molecules with the same activity, making them potentially useful for inducing suicidal germination (*8*).

### Identification of fungal metabolites that can induce seed germination

We first screened a few fungal species owned in our laboratory for fungi that produce compounds that can induce the germination of *Orobanche minor* seeds. We observed that *G. fujikuroi* culture extracts induced the germination of *O. minor* seeds similar to *rac*-GR24, which is a synthetic SL analog (Fig. 1A). Next, on the basis of germination assay results, active compounds were purified from *G. fujikuroi* culture extracts using a series of column chromatography, which yielded the final active fractions Fr. A2A4, Fr. A2A6, Fr. A2A7/A2A8, and Fr. B1B1A (Fig. 1B, figs. S1 to S9). An LC-MS/MS analysis of each active fraction detected several jasmonates, which are plant hormones involved in defense responses, in all active fractions (*9*). To determine the chemical structure of these compounds, we synthesized authentic standards of several jasmonates predicted according to chemical formulae. By comparing the retention time and MS/MS spectrum of purified compounds with those of authentic standards, we determined that Fr. A2A4 contained (−)-jasmonoyl-L-valine ((−)-JA-Val) (fig. S10). Additionally, (+)-4,5-didehydro-jasmonoyl-L-isoleucine ((+)-4,5-ddh-JA-Ile) and its isomers were detected in Fr. A2A6 (fig. S11), whereas (−)-jasmonoyl-L-Ile ((−)-JA-Ile) was detected in Fr. A2A7/A2A8 (fig. S12). A precursor of jasmonic acid (JA), namely *cis*-OPDA, was detected in Fr. B1B1A (fig. S13). Jasmonate production by *G. fujikuroi* and related species has been described in several papers (*10*). To verify the germination-inducing effects of jasmonates on *O. minor* seeds, we performed a germination assay using chemically synthesized or commercially available jasmonates. Assay results showed that in addition to (±)-JA and (±)-methyl jasmonate ((±)-MeJA), which have been reported to induce *O. minor* seed germination (*11*), other jasmonates can induce the germination of *O. minor* seeds (Fig. 1C). Although the potency of jasmonates was substantially lower than that of *rac*-GR24, *cis*-OPDA exhibited the highest germination-inducing activity among the tested jasmonates (Fig. 1C). We also observed that 1,000 μM *cis*-OPDA and (−)-JA-Val weakly, but significantly, induced *Striga hermonthica* seed germination (Fig. 1C). The strong bioactivity of coronatine, a bacterial phytotoxin mimicking the structure of a bioactive jasmonate, (3*R*, 7*S*)-JA-Ile, is associated with the co-receptors CORONATINE INSENSITIVE 1 (COI1) and JASMONATE ZIM DOMAIN (JAZ) (*12*, *13*).

**Fig. 1.**
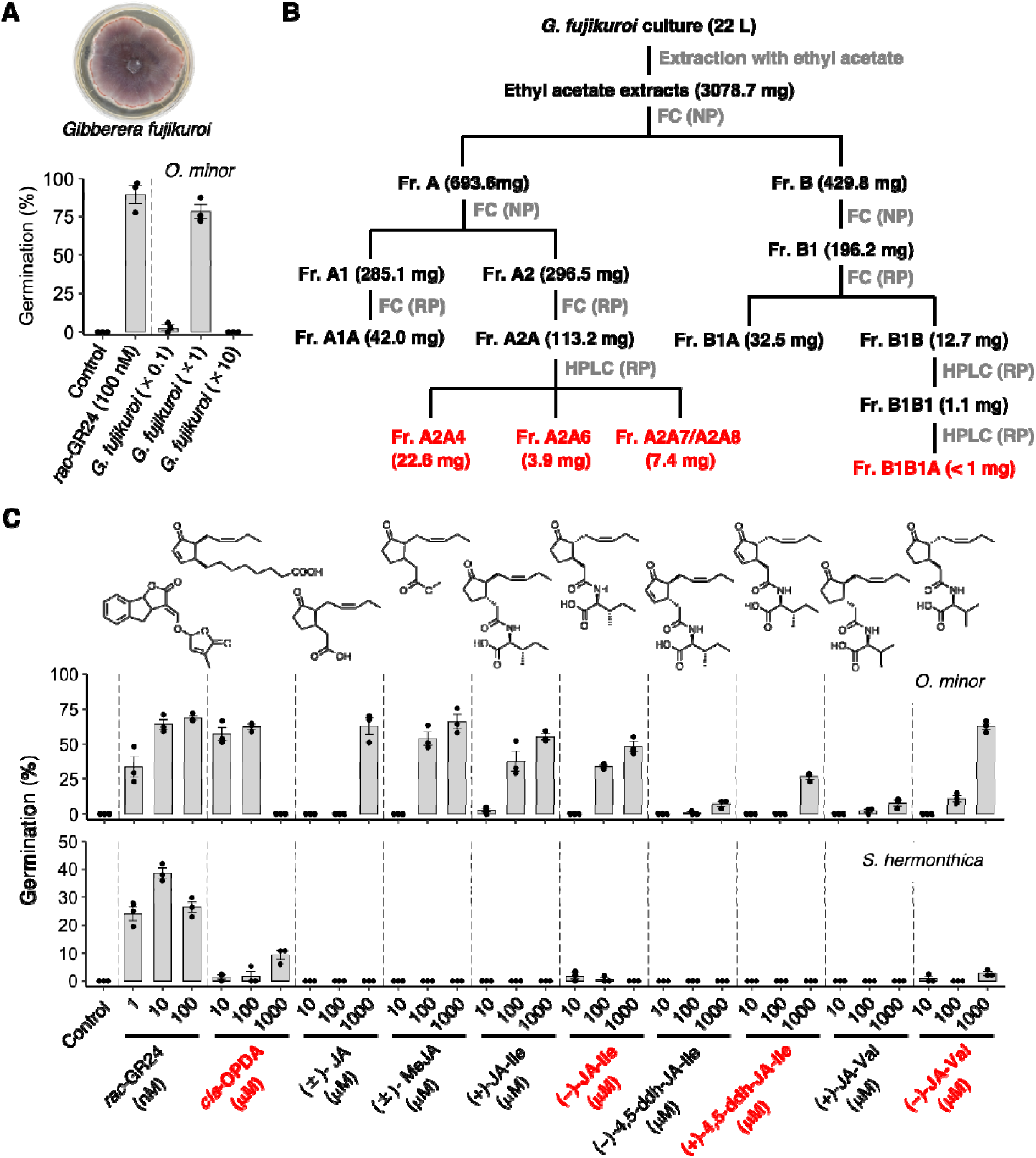
Active compounds in *G. fujikuroi* culture extracts. (**A**) Germination-inducing effects of *G. fujikuroi* culture extracts on *O. minor* seeds. Culture extract concentrations are presented as the fold-enrichment relative to the original culture volume. (**B**) Overview of the purification of functional compounds. Specific details are provided in supplemental materials. Final active fractions are indicated by red text. FC, flash column chromatography; NP, normal phase; RP, reverse phase. (**C**) Germination-inducing effects of chemically or enzymatically synthesized jasmonates on *O. minor* and *S. hermonthica* seeds. Jasmonates in the final active fractions are indicated by red text. Data are presented as the mean ± SE of three biological replicates. Dots represent individual data points.

However, *O. minor* seed germination was weakly induced by coronatine (fig. S14A and B). Considering the relatively weak effect of coronatine, the jasmonate-induced germination of root parasitic plant seeds may be mediated by a pathway other than the typical jasmonate signaling pathway. Jasmonates inhibit seed germination of several plant species in concert with a germination inhibiting plant hormone, abscisic acid (ABA) (*14*, *15*). Conversely, it is intriguing that jasmonates induce root parasitic weed seed germination.

### Synergistic effects of SLs and jasmonates on root parasitic plant seed germination

In some cases, combining two different chemical compounds with the same bioactivity can lead to additive or synergistic effects (*16*). Therefore, we tested whether SLs and jasmonates can synergistically affect *O. minor* seed germination. Notably, when *O. minor* seeds were co-treated with *rac*-GR24 and jasmonates at concentrations that resulted in very low germination rates when these compounds were applied alone, seed germination was strongly induced, reflecting the potential synergistic effects of *rac*-GR24 and jasmonates (Fig. 2A). Similarly, *rac*-GR24 and jasmonates synergistically induced the germination of *S. hermonthica* seeds (Fig. 2A). To quantitatively evaluate these synergistic effects, we performed an *O. minor* seed germination assay using a wide range of jasmonate concentrations with and without 50 pM *rac*-GR24. A low concentration of *rac*-GR24 substantially enhanced the germination-inducing activity of all three jasmonates (Fig. 2B). The EC_50_ values of *cis*-OPDA, (±)-JA, and (−)-JA-Ile decreased significantly when 50 pM *rac*-GR24 was included (*cis*-OPDA: 0.98 μM to 0.011 μM; (±)-JA: >100 μM to 0.22 μM; (−)-JA-Ile: 55.4 μM to 0.72 μM) (Fig. 2B). Similarly, the EC_50_ value of *rac*-GR24 was 200-fold lower in the presence of 100 nM *cis*-OPDA than in the absence of *cis*-OPDA (Fig. 2C). In terms of the utility of jasmonates for enhancing the suicidal germination-inducing effects of SLs, cost-effective and environmentally friendly compounds should be used. Thus, we evaluated the activity of Jasmomate, which is a commercially available plant growth regulator containing propyldihydrojasmonate as the active ingredient. The application of Jasmomate induced germination, but it also increased the effects of *rac*-GR24 on *O. minor* seed germination (Fig. 2D, fig. S15A and B). Similarly, Jasmomate increased the effects of *rac*-GR24 on *S. hermonthica* seed germination, but it had no effect when applied alone (Fig. 2E, fig. S15). Notably, the activity of sphynolactone-7 (SPL-7), which is the most potent inducer of *S. hermonthica* seed germination, was also enhanced by Jasmomate (Fig. 2E) (*17*). These results highlight the effectiveness of jasmonates to amplify the effects of SLs on suicidal germination.

**Fig. 2.**
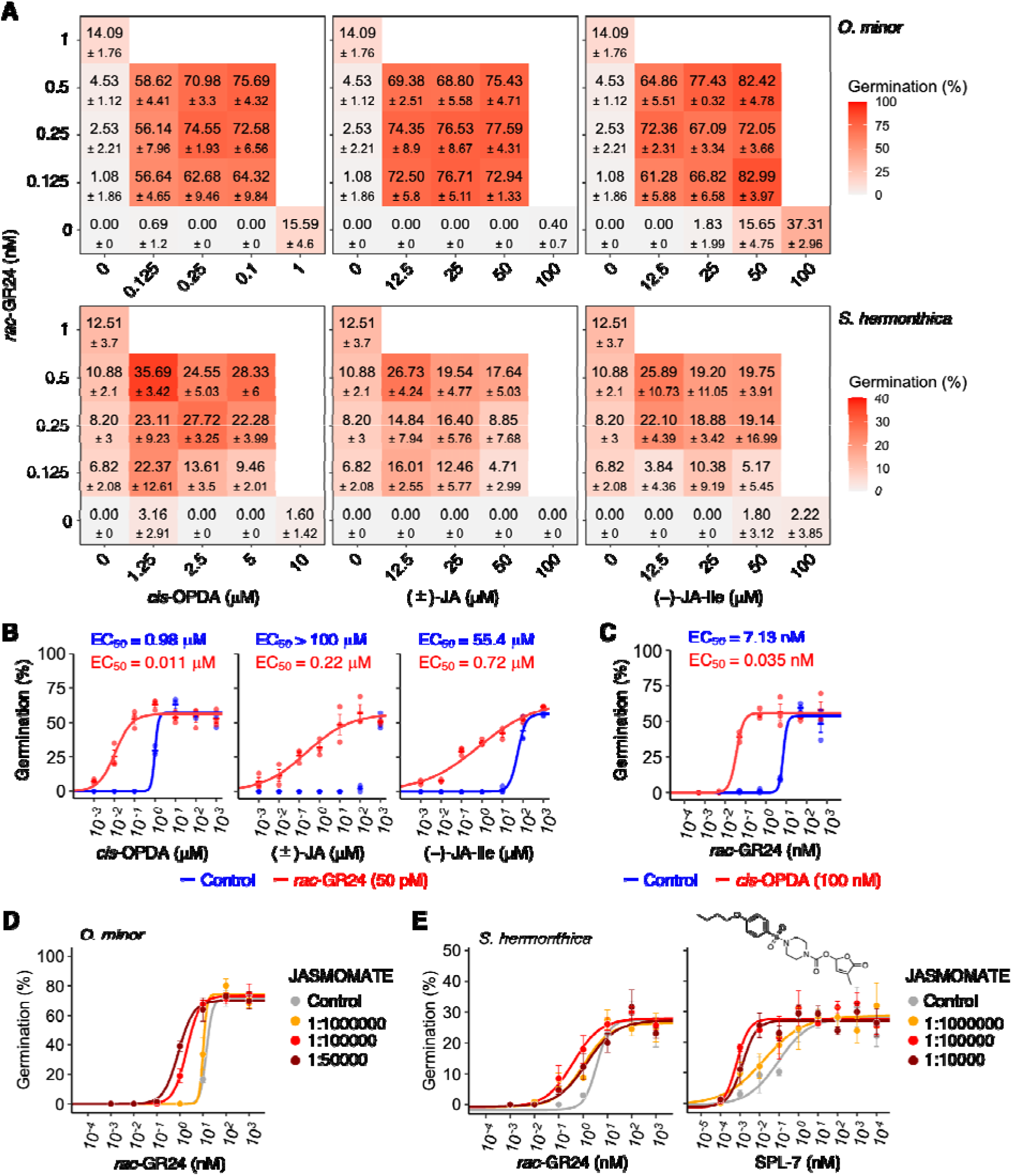
Synergistic effects of jasmonates and SLs on *O. minor* and *S. hermonthica* seed germination. (**A**) Dose-response matrix of three jasmonate–*rac*-GR24 pairs for *O. minor* and *S. hermonthica* seed germination. Data for the *rac*-GR24 treatment alone are provided in each panel for comparison. (**B**) Titration curves of jasmonates in the presence or absence of *rac*-GR24 during *O. minor* seed germination. (**C**) Titration curve of *rac*-GR24 in the presence or absence of *cis*-OPDA during *O. minor* seed germination. (**B, C**) Dots represent individual data points. (**D**) Effect of Jasmomate on the germination-inducing effect of *rac*-GR24 on *O. minor* seeds. (**E**) Effect of Jasmomate on the germination-inducing effect of *rac*-GR24 or SPL-7 on *S. hermonthica* seeds. Data are presented as the mean ± SE of three biological replicates.

### The mechanism of jasmonate perception as a germination signal

SLs were reported to be perceived by DWARF14 (D14) when they act as plant hormones (*18–21*). D14 has a closely related homolog called KARRIKIN INSENSITIVE2 (KAI2), which is an orphan receptor highly conserved in land plants (*22–24*). Root parasitic plants possess a uniquely diverged clade of KAI2, namely KAI2d, some of which have been shown to act as the SL receptors in their seed germination step (*25–29*). In addition to SLs, other host root-derived molecules such as sesquiterpene lactones (STLs) and isothiocyanates (ITCs) were reported to induce root parasitic plants germination and such non-SL type compounds are also reportedly perceived by KAI2d (*28*, *29*). On the basis of the broad range of ligands for KAI2ds and the weak effects of coronatine, we hypothesized that jasmonates induce root parasitic plant seed germination via KAI2d rather than through the COI1–JAZ pathway. According to an earlier genomic analysis, *O. minor* has 11 copies of *KAI2d* (*30*). We selected four KAI2ds (OmKAI2d3/4/9/10) based on phylogenetic diversity and the availability of soluble recombinant proteins. We previously functionally characterized OmKAI2d1–OmKAI2d5, and identified OmKAI2d3/4 as the SL receptor (*29*). We conducted differential scanning fluorimetry and GR24 hydrolysis assays, which indicated that the newly identified OmKAI2d9/10 can bind to and hydrolyze SLs similar to the previously characterized OmKAI2d3/4 (fig. S16A and B) (*29*, *31*). Next, to determine whether *cis*-OPDA can interact directly with OmKAI2d3/4/9/10, fluorescein-labeled *cis*-OPDA (OPDA-PEG3-FL) was synthesized for a fluorescence polarization (FP) assay (Fig. 3A). We confirmed that OPDA-PEG3-FL and *cis*-OPDA have the same germination-inducing activities (Fig. 3B). In the FP assay, binding signals increased when OmKAI2d3/4/9 was incubated with OPDA-PEG3-FL, with *K*_D_ values of approximately 50 μM (Fig. 3C). The binding signals for OmKAI2d3/4/9 clearly decreased when unlabeled *cis*-OPDA was added, confirming the specificity of these interactions (Fig. 3D). The interaction between *cis*-OPDA and OmKAI2d3 was further verified by isothermal titration calorimetry, which revealed a moderate binding affinity (*K*_D_ = 19.8 μM) (fig. S17). FP competition assays also showed that (−)-JA-Ile slightly inhibited the binding of OPDA-PEG3-FL to OmKAI2d3/4/9 (Fig. 3D), consistent with its weak germination-inducing activity. In addition, *rac*-GR24 did not disrupt the binding between OPDA-PEG3-FL and OmKAI2d3/9, but it weakly inhibited the binding to OmKAI2d4 (Fig. 3D). These results suggest that *cis*-OPDA and possibly other jasmonates can bind to OmKAI2d3/9 at unknown allosteric sites.

**Fig. 3.**
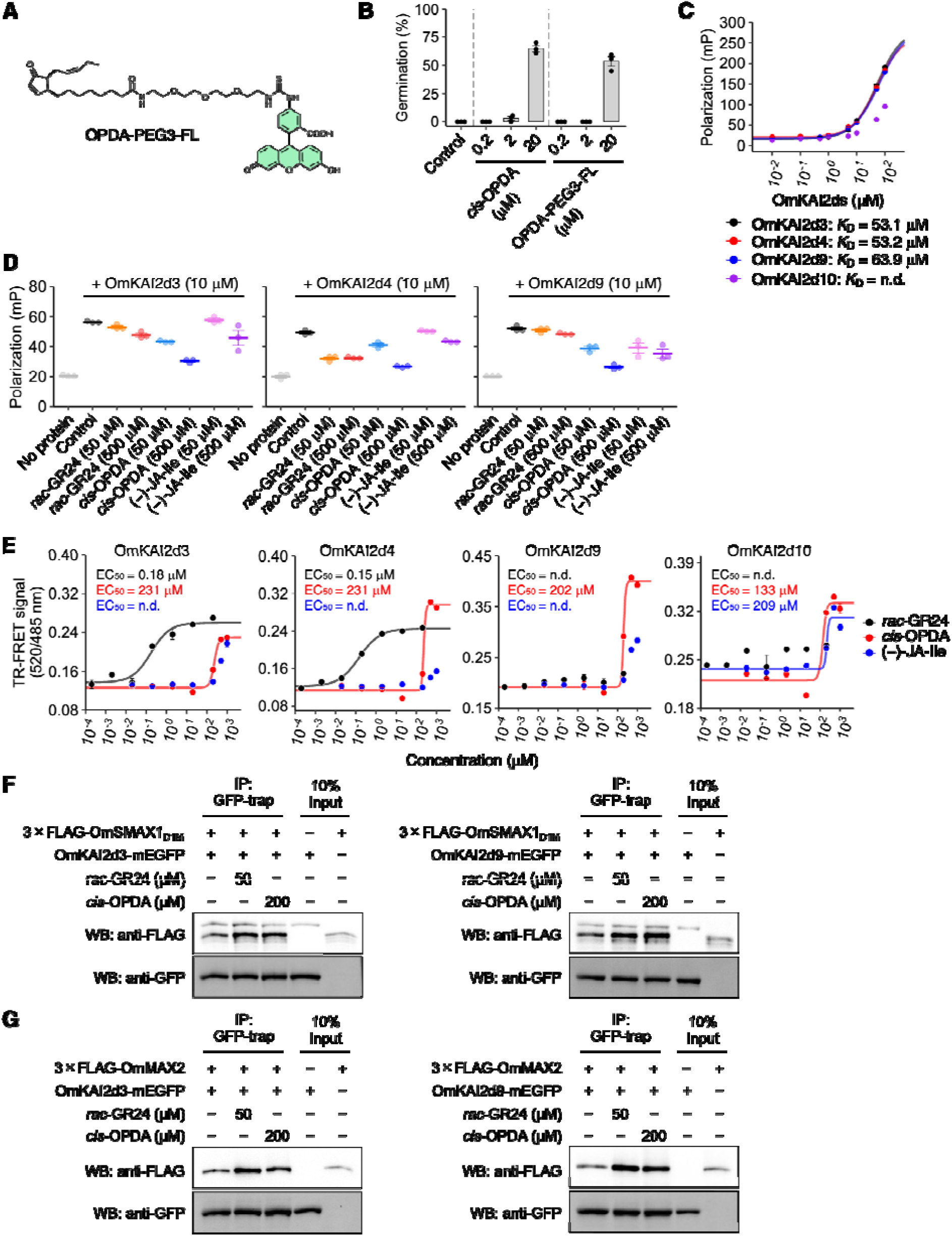
Evaluation of the jasmonate effects on KAI2ds. (**A**) Chemical structure of OPDA-PEG3-FL. (**B**) Comparison of the germination-inducing effects of unlabeled *cis*-OPDA and OPDA-PEG3-FL on *O. minor* seeds. Data are presented as the mean ± SE of three biological replicates. (**C**) Direct interaction between OPDA-PEG3-FL and OmKAI2d proteins as revealed by fluorescence polarization. (**D**) Inhibition of OPDA-PEG3-FL binding to OmKAI2d3/4/9 by *rac*-GR24, *cis*-OPDA, and (−)-JA-Ile. (**E**) Dose-titration of *rac*-GR24, *cis*-OPDA, and (−)-JA-Ile in OmKAI2d3/4/9/10–OmSMAX1_D1M_ TR-FRET assays at a single time point (300 min). (**C, D, E**) Data are presented as the mean ± SE of technical replicates. (**B, D**) Dots represent individual data points. (**F, G**) GFP pull-down assays to detect ligand-induced OmKAI2d3/9–OmSMAX1_D1M_ or OmKAI2d3/9–OmMAX2 interactions. IP, immunoprecipitation; WB, western blot. Uncropped immunoblot images are presented in fig. S23A and B.

During the germination of root parasitic plant seeds, the SL signaling pathway is triggered by the proteasomal degradation of the repressor protein SUPPRESSOR OF MAX2 1 (SMAX1) after a complex comprising SMAX1, KAI2d, and the E3 ubiquitin ligase machinery including an F-box protein, MORE AXILLARY GROWTH (MAX2), is formed (*32*). Thus, we investigated whether jasmonates induce protein–protein interactions (PPIs) involving the SL signaling components. We recently developed a method for monitoring the ligand-induced KAI2d–SMAX1 interaction on the basis of time-resolved Förster resonance energy transfer (TR-FRET) (*33*). We found that *cis*-OPDA and (−)-JA-Ile as well as *rac*-GR24 increased the TR-FRET signals resulting from the interaction between OmKAI2d3/4 and the OmSMAX1-D1M domain (OmSMAX1_D1M_), which is important for the ligand-induced interaction of SMAX1 with KAI2 (Fig. 3E) (*34*). Interestingly, although *rac*-GR24 weakly increased TR-FRET signals resulting from the interaction between OmKAI2d9/10 and OmSMAX1_D1M_, *cis*-OPDA and (−)-JA-Ile significantly increased PPI signals (Fig. 3E). Pull-down assay results reflected the potency of the effects of *cis*-OPDA as well as *rac*-GR24 on the interaction between OmKAI2d3/9 and OmSMAX1_D1M_ (Fig. 3F). We also observed that the OmKAI2d3–OmMAX2 interaction tended to be facilitated by *cis*-OPDA in addition to *rac*-GR24 (Fig. 3G). The interaction between OmKAI2d9 and OmMAX2 was promoted by both *cis*-OPDA and *rac*-GR24 (Fig. 3G). Considered together, these results suggest that jasmonates induce root parasitic plant seed germination by promoting the SL signaling complex formation. Notably, jasmonates and *rac*-GR24 differed in terms of their targeted KAI2d–OmSMAX1_D1M_ complexes under our TR-FRET assay conditions. Thus, combining *rac*-GR24 and jasmonates may activate diverse duplicated KAI2d proteins. This may be the mechanism underlying the synergistic germination-inducing effects of SLs and jasmonates.

### Synergistic activation of OmKAI2ds by SLs and jasmonates

According to FP competition assays, we noticed that *rac*-GR24 progressively increased the signals corresponding to the binding between OPDA-PEG3-FL and all tested OmKAI2d proteins over time (Fig. 4A). We also determined that OmKAI2d3^S95A^, which is a non-functional OmKAI2d3 lacking the catalytic serine residue, was still able to bind to *cis*-OPDA, but impaired the *rac*-GR24-induced increase in FP signals (fig. S18). On the basis of these results and the reactivity of the electrophilic enone moiety of *cis*-OPDA, we hypothesized that OPDA-PEG3-FL covalently binds to OmKAI2d proteins with an altered conformation due to SL binding (*35*, *36*). To test this hypothesis, we performed a non-reducing SDS-PAGE analysis of OmKAI2d3 incubated with OPDA-PEG3-FL in the presence of either the highly active isomer (+)-GR24 or the weakly active isomer (−)-GR24 (*37*). The fluorescence of the band corresponding to OmKAI2d3 modified by OPDA-PEG3-FL increased in the presence of (+)-GR24, but decreased in the presence of unlabeled *cis*-OPDA (Fig. 4B). These results demonstrate that OPDA-PEG3-FL covalently binds to OmKAI2d3 through a process facilitated by (+)-GR24. Consistent with FP assay results, the covalent binding of OPDA-PEG3-FL to OmKAI2d3^S95A^ was not enhanced by (+)-GR24 (Fig. 4B). These observations imply that SL-induced conformational changes to OmKAI2d proteins enable the covalent binding of *cis*-OPDA (Fig. 4C).

**Fig. 4.**
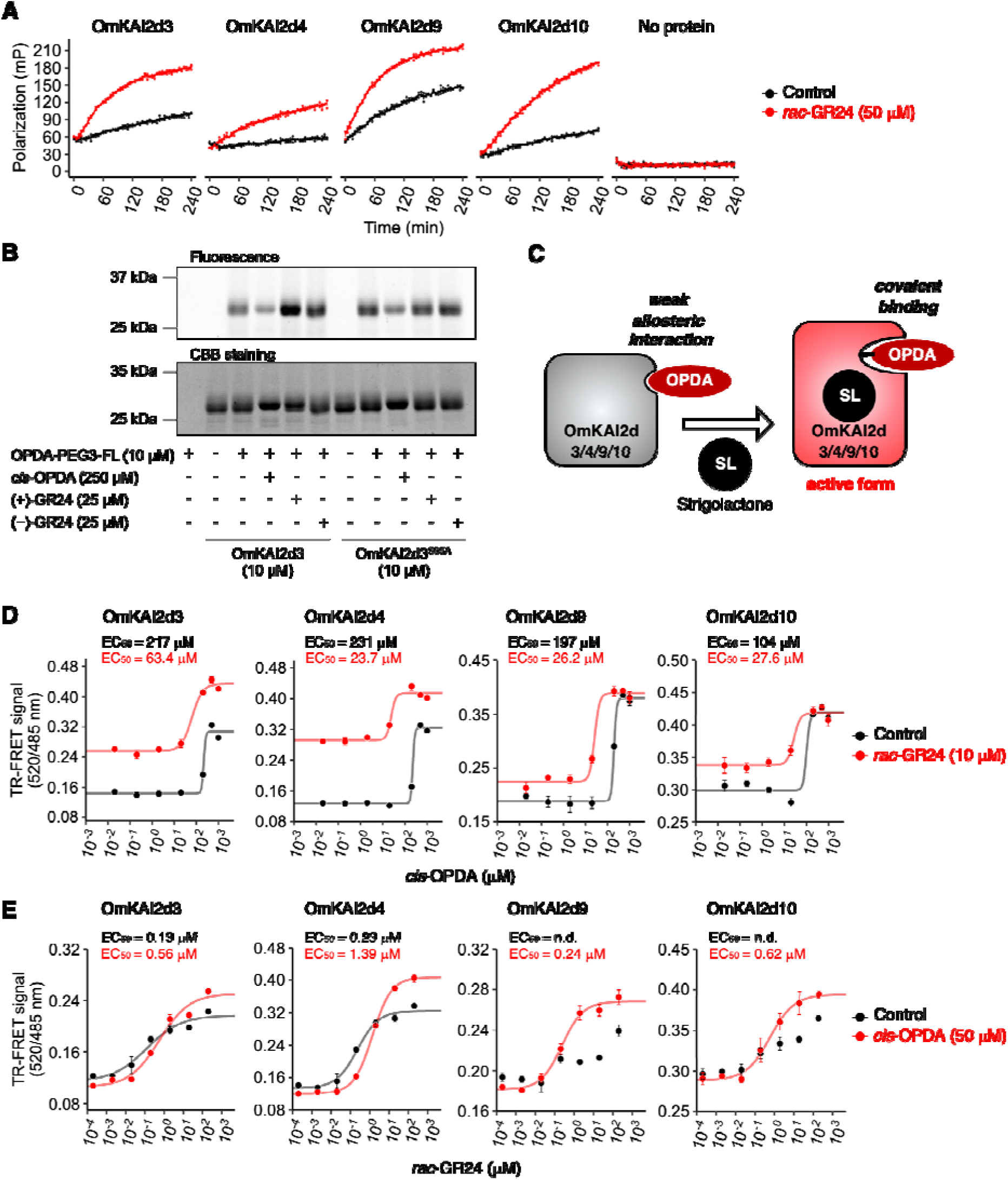
Evaluation of the synergistic effect of jasmonates and SLs *in vitro*. (**A**) Kinetic analyses of OPDA-PEG3-FL binding to OmKAI2d proteins based on fluorescence polarization. Data are presented as the mean ± SE of technical replicates. Dots represent individual data points (**B**) Non-reducing SDS-PAGE analysis of OmKAI2d3 or OmKAI2d3^S95A^ proteins incubated with OPDA-PEG3-FL with or without test chemicals. Fluorescent bands represent OmKAI2d3 covalently bound to OPDA-PEG3-FL. Uncropped gel images are presented in fig. S24. (**C**) Putative model of the SL-enhanced binding of *cis*-OPDA to OmKAI2d proteins. (**D**) Dose-titration of *cis*-OPDA in the presence or absence of *rac*-GR24 in OmKAI2d3/4/9/10–OmSMAX1_D1M_ TR-FRET assays at a single time point (300 min). Some of the data for OmKAI2d9 were extracted from fig. S19. (**E**) Dose-titration of *rac*-GR24 in the presence or absence of *cis*-OPDA in OmKAI2d3/4/9/10–OmSMAX1_D1M_ TR-FRET assays at a single time point (300 min). (**D, E**) Data are presented as the mean ± SE of three technical replicates.

We subsequently investigated whether *rac*-GR24 and *cis*-OPDA cooperatively induce signaling complex formation. In TR-FRET assays involving OmKAI2d3/4/9/10 and OmSMAX1_D1M_, the EC_50_ values of *cis*-OPDA decreased in the presence of 10 µM *rac*-GR24 than in the absence of *rac*-GR24, implying that SLs enhance the effects of *cis*-OPDA on PPIs (Fig. 4D). However, the EC_50_ values of *rac*-GR24 for the formation of the OmKAI2d3/4–OmSMAX1_D1M_ complex increased slightly in the presence of *cis*-OPDA, while *cis*-OPDA increased the maximum values of *rac*-GR24-induced TR-FRET signals (Fig. 4E). Interestingly, the presence of both *rac*-GR24 and *cis*-OPDA significantly increased TR-FRET signals for OmKAI2d9/10–OmSMAX1_D1M_ pairs, resulting in clear sigmoidal curves (OmKAI2d9–OmSMAX1_D1M_: EC_50_ for *rac*-GR24 = 0.24 μM; OmKAI2d10–OmSMAX1_D1M_: EC_50_ for *rac*-GR24 = 0.62 μM), whereas *rac*-GR24 alone modestly increased TR-FRET signals (Fig. 4E). In addition, kinetic analyses indicated that OPDA-induced signals for the OmKAI2d9–OmSMAX1_D1M_ pair increased more quickly in the presence of *rac*-GR24 than in its absence (fig. S19). By contrast, *rac*-GR24 had relatively little effect on the PPI-inducing activity of (−)-JA-Ile (compared with the effects of *cis*-OPDA) (fig. S20A and B). Thus, biochemical assay results suggest that SLs and jasmonates activate OmKAI2ds through different binding sites and eventually induce the formation of the OmKAI2ds–OmSMAX1 complex in a synergistic manner. The cooperative activation of individual OmKAI2ds likely contributes to the synergistic germination-inducing effects of SLs and jasmonates.

### Enhanced sensitivity of root parasitic plants to SL due to host root-derived jasmonates

Given that jasmonates, especially MeJA, act as signaling molecules that mediate plant interactions with other plants and microbes, we hypothesized that root parasitic plants may use root-derived jasmonates as signals for enhancing the sensitivity to SLs during germination (*38*, *39*). However, little is known about the exudation of non-volatile jasmonates, including *cis*-OPDA, JA, and JA-Ile (*40*). Considering jasmonate biosynthesis is induced by mechanical wounding or microbial elicitors, jasmonates are expected to be secreted from roots into the rhizosphere under such conditions (*41*). Thus, we examined whether treating rice plants in a hydroponic culture system with chitin oligosaccharides (COs) can trigger the exudation of non-volatile jasmonates from the roots. A COs treatment significantly promoted the exudation of *cis*-OPDA, JA, and JA-Ile in a CHITIN ELICITOR RECEPTOR KINASE1 (CERK1)-dependent manner (Fig. 5A, B) (*42*, *43*). Next, we analyzed the effects of the root exudates of treated rice plants on *O. minor* seed germination. To specifically examine the enhancer activity of root exudates (i.e., not the germination-inducing effects of SLs in root exudates), we used the SL-deficient mutant *dwarf 10* (*d10*) grown under phosphate-sufficient conditions (*4*, *44*). The results indicated that root exudates from treated *d10* plants enhanced *rac*-GR24-induced germination more effectively than root exudates from untreated plants; these findings were in accordance with jasmonate levels (Fig. 5B, fig. S21A). Accordingly, COs increased the amount of SL enhancers produced by *d10*. To assess the contribution of jasmonates, we compared SL enhancer activities between wild-type plants and a *coleoptile photomorphogenesis 2* (*cpm2*) mutant lacking ALLENE OXIDE CYCLASE, the key enzyme for *cis*-OPDA formation (*45*). Consistent with the impaired jasmonate biosynthesis of *cpm2*, SL enhancer activities and related elicitor effects were undetectable in *cpm2* root exudates, but they were observed in wild-type root exudates (Fig. 5C, fig. S21B). These results suggest that root parasitic plants exploit elicitor-induced jasmonates exuded from host roots to enhance SL activities during the host recognition process (Fig. 5D).

**Fig. 5.**
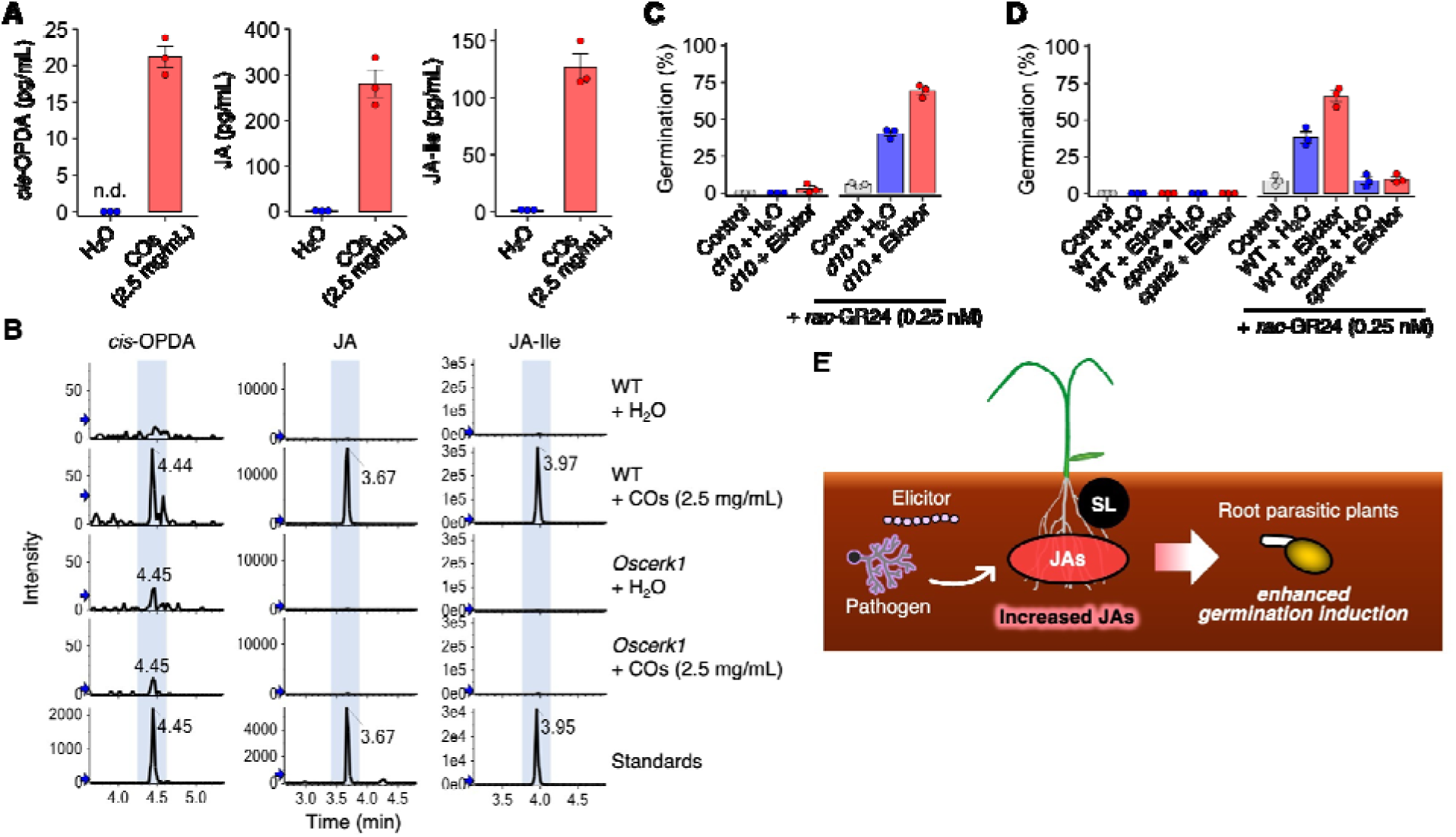
SL enhancer effects of jasmonates in host root exudates. (**A**) LC-MS/MS analysis of *cis*-OPDA, JA, and JA-Ile in rice (BL2) hydroponic cultures after a 2-h treatment with chitin oligosaccharides (COs). Data are presented as the mean ± SE of three biological replicates. (**B**) OsCERK1-dependent elicitor-induced exudation of jasmonates. Jasmonates in hydroponic cultures of BL2 wild-type and *Oscerk1* plants treated with COs for 2 h were analyzed by LC-MS/MS. LC-MS/MS chromatograms of *cis*-OPDA, JA, JA-Ile, and the corresponding internal standards in the multiple reaction monitoring mode are presented. (**C, D**) SL enhancer effects of rice root exudates obtained after a 2-h treatment with 2.5 mg/mL COs on *O. minor* seed germination. *O. minor* seeds were co-treated with ethyl acetate extracts of root exudates (20-fold concentrated) and *rac*-GR24. Control seeds were not treated with root exudate extracts. Data are presented as the mean ± SE of three biological replicates. (**E**) Putative model of the biological significance of the enhanced sensitivity of root parasitic plant seeds to SL due to host root-derived jasmonates.

## Discussion

We determined that jasmonates stimulate root parasitic plant seed germination by activating KAI2d proteins, possibly reflecting the promiscuity of these receptors. Recent studies demonstrated that D14 can recognize some compounds with structures that differ from those of SLs, including cyclodipeptide (*7*) and zaxinone (*46*). Moreover, considering that KAI2d can interact with ITCs or STLs, the observed promiscuity might be a common characteristic of KAI2d/D14 receptor family members. Such a feature of parasitic plant KAI2d proteins may be conducive to effective host recognition because it enables the detection of SLs as well as other host-derived compounds. This ligand diversification may be enhanced by receptor duplications in root parasitic plants.

We also observed that jasmonates and SLs can synergistically induce *O. minor* and *S. hermonthica* seed germination. On the basis of biochemical data, we propose the following two mechanisms underlying the synergistic effects of jasmonates and SLs: (i) activation of diverse and highly duplicated KAI2d proteins; (ii) synergistic activation of individual OmKAI2d proteins through an allosteric mechanism (fig. S22). To the best of our knowledge, this is the first report on the allosteric regulatory system of plant hormone receptors for two different hormones.

Combining chemical treatments is a promising strategy to increase efficacy, thereby preventing the emergence of resistant organisms, while also decreasing the side effects of the applied chemicals (*16*). Earlier attempts to develop highly potent SL analogs resulted in the identification of SPL-7 as a suicidal germination inducer, even at femtomolar concentrations; however, methods involving combined chemical treatments have not been considered (*17*). Notably, our study demonstrated the effectiveness of combining SLs and jasmonates to synergistically induce *O. minor* and *S. hermonthica* seed germination. As mentioned above, jasmonates were produced by several fungi at levels greater than those generated by plants. Thus, fermentation-based jasmonate production using such fungi may be relevant to the large-scale preparation of SL enhancers. Furthermore, we demonstrated that a commercially available agrochemical, Jasmomate, can increase the efficacy of *rac*-GR24 and SPL-7. The strong synergism between these two phytohormones may be exploited to develop an effective method for eliminating root parasitic plants.

The exudation of SLs from roots is strongly enhanced under nutrient-deficient conditions (*47*). The seed germination of *Phelipanche ramosa*, an obligate root parasitic plant, is induced by SLs as well as *Brassica*-specific ITCs biosynthesized as defense-related compounds in response to herbivore attack (*48*). Furthermore, we revealed that elicitor-induced jasmonates secreted by rice roots can increase the sensitivity of *O. minor* seeds to SL. Accordingly, by detecting stress-induced chemicals, root parasitic plants may recognize plants under such stress conditions as better hosts than healthy plants. However, additional studies are required to determine whether root parasitic plants benefit from targeting stressed plants. Our findings suggest that the duplication of *KAI2d* genes enabled root parasitic plants to recognize molecules other than SLs possibly by increasing ligand diversity. The host root secretes various chemicals into the rhizosphere, but root exudate chemical compositions fluctuate in response to environmental factors. The synergistic effects of SLs and jasmonates imply that root parasitic plants may increase the likelihood they detect a suitable host by sensing SLs as well as other chemicals using highly duplicated and promiscuous KAI2d receptors.

## Acknowledgments

We thank Nagasawa Water Purification Plant for the corporation regarding *O*. *minor* sampling. We thank Dr. Steven Runo for kindly providing seeds of *S. hermonthica*. We thank Edanz for editing a draft of this manuscript.

## Funding

Japan Society for the Promotion of Science KAKENHI grants 19K05852 (YS), 22H02276 (YS), 23H05409 (YS), 24H00878 (YS) and 24KJ2051 (TS)

JST FOREST Program grant JPMJFR211S (YS) JST ACT-X grant JPMJAX22BH (KN)

Mitsubishi Foundation (YS)

Kato Memorial Bioscience Foundation (YS).

## Author contributions

Conceptualization: YS

Methodology: TS, TI, MB, MO, HK, KN, YS Investigation: TS, TI, MB, MO, JW, MK Visualization: TS

Funding acquisition: TS, KN, YS Project administration: YS Supervision: KN, YS

Writing – original draft: TS, KN, YS

Writing – review & editing: TS, MB, MO, KM, HK, HM, KN, YS

## Diversity, equity, ethics, and inclusion [optional]

We welcome statements pertaining to diversity, equity, ethics, and inclusion, as outlined in our Editorial Policies.

## Competing interests

Authors declare that they have no competing interests.

## Data and materials availability

All data are available in the main text or the supplementary materials.

## Supplementary Materials

Materials and Methods

Figs. S1 to S27

Tables S1 to S3

References (*49–62*)

## Notes

### Competing Interest Statement

The authors have declared no competing interest.

### Summary of Updates

There were typos in the first author name. We corrected them.

