## Supplementary file for "Two phytohormones synergistically induce parasitic weeds seed germination via KAI2d receptors"

### Materials and Methods

#### General methods

Flash column chromatography were performed by Smart Flash W-Prep 2XY (Yamazen) equipped with a disposable column, Universal Column Premium silica gel or Universal Column ODS (Yamazen). NMR spectra were measured by a NMR spectrometer JNM-ECZL, JNM-AL300 (JEOL), or JNM-ECZ 400. High-resolution mass spectrometry (HRMS) was performed by a quadrupole/time-of-flight tandem mass spectrometer X500R (AB SCIEX) or a JMS-T100GCV (JEOL) instrument. All reagents for chemical syntheses were purchased from Tokyo Chemical Industry Co., Ltd. (Tokyo, Japan) and FUJIFILM Wako Pure Chemical Corp. (Osaka, Japan).

#### Chemicals

(+)-*cis*-OPDA was synthesized according to the previous paper, with slight modification (*49*). We used *Arabidopsis thaliana* allene oxide cyclase 2 (AtAOC2), prepared as previously described (*50*), for the enzymatic synthesis of (+)-*cis*-OPDA. (+/−)-JA-Ile, (+/−)-ddh-JA-Ile, (+/−)-JA-Val were synthesized as described in previous papers (*51*, *52*). (+)-GR24 and (−)-GR24 were separated from commercially available *rac*-GR24 (Chiralix) as previously described (*33*). Coronatine was purchased from Sigma Aldrich. JASMOMATE was purchased from Mitsui Chemicals Crop & Life Solutions, Inc. JA *d*6, JA-Ile *d*6 were prepared as previously described (*53*). *trans*-OPDA *d*6 was prepared according to a previously reported method using propyltriphenylphosphodium iodide-*d*7 instead of propyltriphenylphosphonium iodide (*54*). The $^{1}$H NMR, $^{13}$C NMR and HRMS data of the *trans*-OPDA *d*6 were recorded as follow. $^{1}$H NMR (400 MHz, $CDCl_3$) δ 7.58 (1 H, dd, *J*=5.8, 2.6 Hz), 6.10 (1H, dd, *J*=5.8, 1.8 Hz), 5.24 (1H, tt, *J*=7.2, 4.8 Hz), 2.55 (1H, m), 2.43 (1H, ddd, *J*=14.8, 7.6, 4.8 Hz), 2.34 (2H, t, J=7.6), 2.28 (1H, m), 1.98 (1H, ddd, *J*=8.0, 4.8, 2.4 Hz), 1.61 (2H, m), 1.38-1.23 (8H m). $^{13}$C NMR (100 MHz, $CDCl_3$) δ 211.95, 179.15, 167.64, 132.92, 125.03, 51.5, 47.13, 34.35, 34.35, 33.94, 29.78, 29.14, 29.02, 27.46, 24.67 ppm; HRMS (FD): *m*/*z* calculated for $C_{18}H_{22}D_6O_3$, : 298.24150$[M]^+$; found: 298.24091.

#### Synthesis of OPDA-PEG3-FL

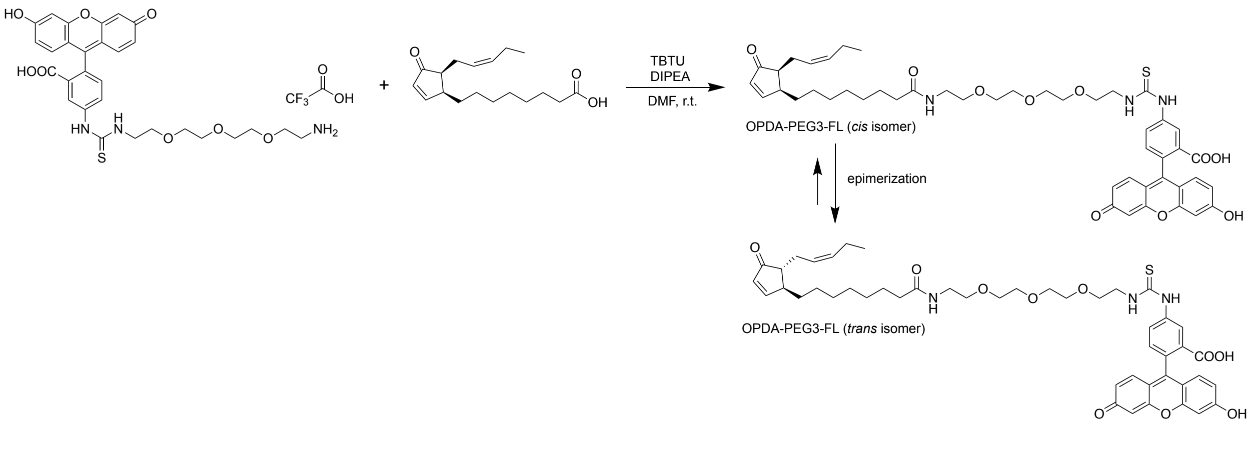

*tert*-Butyl (2-(2-(2-(2-aminoethoxy)ethoxy)ethoxy)ethyl)carbamate (0.087 mmol), fluorescein 5-isothiocyanate (0.081 mmol), *N*,*N*-diisopropylethylamine (DIPEA) (0.20 mmol) and *N*,*N*-dimethylformamide (DMF) (1.0 mL) were added to a flask containing a magnetic stirring bar. After stirring the mixture at room temperature for 1 hour, water was added. The solution was concentrated *in vacuo*. The crude sample was purified by flash column chromatography on ODS using gradient elution with acetonitrile in water (0–2 min: 5% acetonitrile, 3.01–12.0 min: 5–70%) to obtain Boc-protected PEG3-FL. The orange solid was dissolved in trifluoroacetic acid (TFA) (0.5 mL). After incubation for 2 hours, the solution was dried to afford PEG3-FL as TFA salt (54.2 mg, 0.078 mmol, 96%). The product was directly used to synthesize OPDA-PEG3-FL.

(+)-*cis*-OPDA (0.036 mmol), PEG3-FL (0.03 mmol), *N*,*N*-diisopropylethylamine (DIPEA) (0.11 mmol), 2-(1*H*-benzo[*d*][1,2,3]triazol-1-yl)-1,1,3,3-tetramethylisouronium tetrafluoroborate (TBTU) (0.036 mmol) and *N*,*N*-dimethylformamide (DMF) (0.12 mL) were added to a flask containing a magnetic stirring bar. After stirring the mixture at room temperature for 1 day, 1N HCl was added. The precipitate was collected and washed with water. The sample was purified by flash column chromatography on ODS using gradient elution with methanol in water (0–3 min: 5% methanol, 3.01–18.0 min: 5–85%, 18.01–25.0 min: 85%) to afford the *cis*/*trans* mixture of OPDA-PEG3-FL (10.4 mg, 0.012 mmol, 33% yield). The *trans* isomer was presumed to arise from the *cis* isomer by epimerization during the reaction and/or sample preparation process (*55*). The ratio of *cis* to *trans* isomers was determined to be approximately 4 : 1 by $^{1}$H NMR spectroscopy (δ = 6.14 (dd, *J* = 6.0, 2.0 Hz, 1H) for *cis* isomer, δ = 6.09 (dd, *J* = 6.0, 2.0 Hz, 1H) for *trans* isomer (fig. S25). The *cis*/*trans* mixture of OPDA-PEG3-FL was use for the experiments without further purification. The $^{1}$H NMR (fig. S25), $^{13}$C NMR (fig. S26) and HRMS data of the *cis* isomer were recorded as follow. $^{1}$H NMR (500 MHz, $CDCl_3$) δ 8.15 (s, 1 H), 7.89 (dd, *J* = 6.0, 3.0 Hz, 1H), 7.77 (d, *J* = 5.5 Hz, 1H), 7.19–7.17 (m, 1H), 6.94 (d, *J* = 6.5 Hz, 2H), 6.65–6.64 (m, 2H), 6.60–6.57 (m, 2H), 6.14 (dd, *J* = 6.0, 2.0 Hz, 1H), 5.43–5.32 (m, 2H), 3.83 (br, 2H), 3.73–3.57 (m, 12H), 3.50 (t, *J* = 5.0 Hz, 2H), 3.01 (br, 1H), 2.49–2.39 (m, 2H), 2.17–2.14 (m, 3H), 2.07–2.01 (m, 2H), 1.76–1.70 (m, 1H), 1.42–1.24 (br, 8H), 1.18–1.11 (m, H), 0.95 (t, *J* = 7.5 Hz, 3H); $^{13}$C NMR (125 MHz, $CDCl_3$) δ 14.24, 21.76, 24.94, 26.99, 28.64, 30.22, 30.31, 30.78, 31.82, 37.05, 40.32, 45.48, 45.81, 50.99, 70.21, 70.61, 71.26, 71.32, 71.58, 71.63, 103.82, 113.81, 126.09, 128.28, 131.64, 132.82, 133.29, 133.75, 134.92, 141.99, 157.00, 170.32, 170.67, 172.85, 176.39, 182.80, 213.61 ppm; HRMS (ESI): *m*/*z* calculated for $C_{47}H_{57}N_3O_{10}S+H^+$: 856.3837 $[M+H]^+$; found: 856.3839.

Note: The peaks of impurities and/or *trans* isomer overlapped around 0.9, 1.3 and 2.1 ppm in the 1H-NMR spectrum chart (fig. S25). Several carbon peaks of the fluorescein moiety were not detected in the $^{13}$C NMR analysis because of its low solubility (fig. S26). The *cis* geometry of side chains at cyclopentenone ring was confirmed by Nuclear Overhauser Effect (NOE) correlations (fig. S27, A to C).

Purification and identification of the active molecules from culture extracts of *G. fujikuroi*

*G. fujikuori* (MAFF No. 241712) was grown on potato dextrose agar (PDA) media at 25°C under dark conditions. Mycelium was transferred into liquid 10% ICI medium (*56*) in Erlenmeyer flask and incubated with shaking at 130 rpm at 25°C under dark conditions for 5-7 days. Culture media was filtrated through four layers of Miracloth (Merck). The filtrate was extracted with ethyl acetate twice. The organic phase was dried over $Na_2SO_4$ and concentrated *in vacuo*. We purified the active compounds from crude extracts (3078.7 mg) obtained from 22 L of *G. fujikuori* culture medium. Crude extracts were fractionated by flash column chromatography on silica gel using mobile phase A (*n*-hexane) and mobile phase B (EtOAc) under the following elution conditions (0–2 min: 30% B, 2.01–14.0 min: 30–100% B, 14.01–25.0 min 100% B). The fractions (Fr. A: 9.0–13.0 min, Fr. B: 14.0–16 min) were pooled, based on the results of *O. minor* germination assay. Fr. A was fractionated by flash column chromatography on silica gel using mobile phase A (*n*-hexane) and mobile phase B (EtOAc) under the following elution conditions (0–2.0 min: 21% B, 2.01–11.5 min: 21–50% B, 11.51–13.0 min: 50% B, 13.01–29.5 min: 50–70% B, 29.51–36.0 min: 70% B, 36.01–42.0 min: 100% B. The fractions (Fr. A1: 9.5–18.5 min, Fr. A2: 21.5–31.5 min) were pooled, based on the results of *O. minor* germination assay. Fr. A1 was fractionated by flash column chromatography on ODS using mobile phase A ($H_2O$ (0.5% AcOH)) and mobile phase B ($CH_3CN$ (0.5% AcOH)) under the following elution conditions (0–3.0 min: 20% B, 3.01–15.0 min: 20–60% B, 15.01–18.0 min: 60% B, 18.01–25.0 min: 100% B). The fractions (Fr. A1A: 12.0–22.0 min) were pooled, based on the results of *O. minor* germination assay. Fr. A2 was fractionated by flash column chromatography on ODS using mobile phase A ($H_2O$ (0.5% AcOH)) and mobile phase B ($CH_3CN$ (0.5% AcOH)) under the following elution conditions (0–3.0 min: 20% B, 3.01–15.0 min: 20–60% B, 15.01–18.0 min: 60% B, 18.01–25.0 min: 100% B). The fractions (Fr. A2A: 12.0–18.0 min) were pooled, based on the results of *O. minor* germination assay. Fr. A2A was fractionated by HPLC equipped with an ODS column (PEGASIL ODS SP100, ϕ 4.6 mm × 250 mm; Senshu Scientific) using mobile phase A ($H_2O$ (0.1% AcOH)) and mobile phase B ($CH_3CN$ (0.1% AcOH)) under the following elution conditions (0–25.0 min: 40–100% B, 25.01–35.0 min: 100% B, 35.01–50.0 min: 40% B). The fractions (Fr. A2A1–A2A8) were pooled as shown in fig. S5, based on the results of *O. minor* germination assay and the HPLC-UV chromatogram. Fr. A2A7 and Fr. A2A8 were combined into a single pool. Fr. B was fractionated by flash column chromatography on silica gel using mobile phase A (*n*-hexane) and mobile phase B (EtOAc) under the following elution conditions (0–3.0 min: 52% B, 3.01–13.0 min: 52–73% B, 13.01–16.0 min: 73% B, 16.01–29.0 min: 73–97% B, 29.01–36.0 min: 97% B). The fractions (Fr. B1: 9.5–12.5 min) were pooled, based on the results of *O. minor* germination assay. Fr. B1 was fractionated by flash column chromatography on ODS using mobile phase A ($H_2O$ (0.5% AcOH)) and mobile phase B ($CH_3CN$ (0.5% AcOH)) under the following elution conditions (0–3.0 min: 20% B, 3.01–15.0 min: 20–60% B, 15.01–18.0 min: 60% B, 18.01–25.0 min: 100% B). The fractions (Fr. B1A: 12.0–17.0 min, Fr. B1B: 17.01–24.0 min) were pooled, based on the results of *O. minor* germination assay. Fr. B1B was fractionated by HPLC equipped with an ODS column (PEGASIL ODS SP100, ϕ 4.6 mm × 250 mm; Senshu Scientific) using mobile phase A ($H_2O$ (0.1% AcOH)) and mobile phase B ($CH_3CN$ (0.1% AcOH)) under the following elution conditions (0–25.0 min: 40–100% B, 25.01–35.0 min: 100% B, 35.01–50.0 min: 40% B). The fractions (Fr. B1B1: 29.0–33.0 min) were pooled, based on the results of *O. minor* germination assay. Fr. B1B1 was fractionated by HPLC equipped with an ODS column (PEGASIL ODS SP100, ϕ 4.6 mm × 250 mm; Senshu Scientific) using mobile phase A ($H_2O$ (0.1% AcOH)) and mobile phase B ($CH_3CN$ (0.1% AcOH)) under the following elution conditions (0–25.0 min: 60% B, 25.01–30.0 min: 80% B, 30.01–45.0 min: 60% B). The fractions (Fr. B1B1A: 26.0–28.0 min, Fr. B1B1B: 28.01–29.0 min) were pooled, based on the results of *O. minor* germination assay. The active final fractions were analyzed by LC-MS/MS equipped with the reverse-phase column (CORTECS UPLC Phenyl, 1.6 μm, ϕ 2.1 × 75 mm; Waters) in information-dependent acquisition (IDA) mode. The obtained data was analyzed by MS-DIAL4 (*9*). The structure of the active compounds was identified by comparing their retention times and MS/MS spectrum, obtained in MRM mode, with those of authentic standards. Detailed information about the analytical condition was described in table S1.

Germination assay of root parasitic plant seeds

Germination assays were performed as previously described, with slight modifications (*57*). Sterilized seeds of *O. minor* and *S. hermonthica* were resuspended in 0.1% agar solution and loaded onto 5 mm glass fiber filter disks (20–100 seeds/disk) on paper filter wetted with sterilized water. *O. minor* seeds were conditioned at 23℃ for 13–15 days. *S. hermonthica* seeds were conditioned at 30℃ for 7–14 days. After conditioning, disks were transferred to clear 96-well plates (AS ONE). A 30 μL aliquot of chemical solutions was added to each well. 96-well plates were incubated for the same conditions as the conditioning. After 4–5 days of chemical treatment, germination rates of *O. minor* seeds were measured. After 1–2 days of chemical treatment, germination rates of *S. hermonthica* seeds were measured.

Preparation of recombinant proteins

The cording sequence of OmKAI2d9 (KAL6535241.1) and OmKAI2d10 (KAL6535305.1) were amplified from cDNA of *O. minor* by PCR with primers (OmKAI2d9-F: 5′-CACCATGAACCGTATAGTTGGACTTG, OmKAI2d9-R: 5′-TCATACATCAACAATATCACTATTTATATGCCGGA, OmKAI2d10-F: 5′-CACCATGAGTAGCATAGTTGGTGC, OmKAI2d10-R: 5′-TTAAATATCGTGATTTATATGCCGGAGCAG). Each DNA fragment was sub-cloned into pENTR D-TOPO using TOPO cloning Kit (Thermo Fisher Scientific). The cording sequence of OmKAI2d9 and OmKAI2d10 were amplified from the sub-cloned vector by PCR with primers (OmKAI2d9-pET-IF-F: 5′-CTCTTTCAGGGACCCATGAACCGTATAGTTGGACT, OmKAI2d9-pET-IF-R: 5′-CGGATCCTGGTACCCTCATACATCAACAATATCACTA, OmKAI2d10-pET-IF-F: 5′-CTCTTTCAGGGACCCATGAGTAGCATAGTTGGTGC, OmKAI2d10-pET-IF-R: 5′-CGGATCCTGGTACCCTTAAATATCGTGATTTATATGCC). Each DNA fragment was cloned into pET-47b (+) by In-Fusion cloning. The protein expression vector for OmKAI2d3$^{S95A}$ was prepared using KOD-Plus-Mutagenesis Kit (TOYOBO) with primers (OmKAI2d3_S95A-F: 5′- GCTCTTTCTTGCATGGCTGCTGC-3′, OmKAI2d3_S95A-R: 5′- GTGTCCAACGTAAATACATTTTCCAGAAC-3′). 6×His-tagged OmKAI2d proteins, GFP-fused OmKAI2d proteins and 3×FLAG-tagged OmSMAX1$_{D1M}$ were prepared as previously described (*33*).

For the construction of 6×His-3×FLAG-OmMAX2 protein expression vector, the insert DNA fragment cording 6×His-3×FLAG-HRV3C sequence was prepared by primer extension using partially complemental oligos (6×His-3×FLAG-HRV3C-F: 5′-CATCGGGCGCGGATCCATGCACCACCACCACCACCACGGTTCCGACTACAAGGACCACGACGGTGACTACAAGGACCACGACATCGACTACAAGGACGACGACGACAAGC-3′, 6×His-3×FLAG-HRV3C-R: 5′-GTAGGCCTTTGAATTCAGGACCCTGGAACAGCACCTCCAGCTTGTCGTCGTCGTCCTTGTAGTCGATGTCGTGGTCCTTGTAGTCACCGTCGTGGTCCTTGTAGTCGGAA-3′). The elongated DNA fragment was cloned into pFastBacDual by In-Fusion cloning.

The cording sequence of OmSKP1(KAL6504770.1) and OmMAX2 (KAL6553804.1) were amplified from cDNA of *O. minor* by PCR with primers (OmSKP1-F: 5′-AAGACTTGATCACCCGGGATGTCGTCATCCGCAGCC-3′, OmSKP1-R: 5′-ATGGCTCGAGATCCCGGGTCACTCAAACGCCCACTGGTTTT-3′, OmMAX2-F: 5′-

CCAGGGTCCTGAATTCATGGCTGTGGCCTCTACTAC-3′, OmMAX2-R: 5′-ACTTCTCGACAAGCTTTCAGTCAGAGATTTGGCGCCC-3′). Each DNA fragment was cloned into the modified pFastBacDual vector by stepwise In-Fusion cloning. 6×His-3×FLAG-OmMAX2 and OmSKP1 were co-expressed by using Bac-to-bac baculovirus expression system with ExpiSf9 cells (Invitrogen) according to the manufacture's protocol. Cells were harvested after 72 h of cultivation and stored at −80°C. Cells from 100 mL cell culture were resuspended in 10 mL of extraction buffer [50 mM HEPES-NaOH (pH 8.0), 150 mL NaCl, 5 mM 2-mercaptoethanol, 50 μM MG-132, 0.5% triton X-100, 10% glycerol, 1× EDTA-free protease inhibitor cocktail (nacali tesque)]. Resuspended cells were lysed with gentle sonication. After centrifugation at 18,000 × g for 30 min, the supernatant was subjected affinity chromatography using TALON Superflow resin (Cytiva). The column was washed with the washing buffer [50 mM HEPES-NaOH (pH 8.0), 150 mM NaCl, 5 mM imidazole, 0.5% triton X-100]. The bound protein was eluted with the elution buffer [1×PBS (−), 150 mM imidazole]. The eluate was concentrated using VIVASPIN Turvo15 (Sartorius) and buffer-exchanged into 1×PBS (–). The protein solution was aliquoted to the appropriate volume, immediately frozen in liquid nitrogen, and stored at −80°C until use.

Differential scanning fluorimetry

OmKAI2d9/10 proteins (5 μg) were dissolved in 20 μL of 1× PBS (–) buffer containing SYPRO Orange (Sigma Aldrich) and each SL analog with 5% (v/v) acetone on a 96-well PCR white plate (Roche). SYPRO Orange (×5000) was diluted to ×1.25 final concentration in a reaction mixture. The mixtures were heated from 20°C to 95°C with scanning the fluorescence intensity (Ex/Em: 483/610 nm) using LightCycler480 (Roche). The first derivative curve was calculated from a plot of the fluorescence intensity against the temperature using the LightCycler480 Software.

GR24 hydrolysis assay

Hydrolysis assays were performed using 7.5 μg of recombinant 6×His-OmKAI2d9/10 proteins in 50 μL of 50 mM phosphate-Na buffer (pH 7.0) containing 10 μM each (+)-GR24 with 1% (v/v) acetone. After incubation at 30ºC for 1 hour, the enzymatic reaction was quenched by the addition of nine equal volumes of acetonitrile containing 40 ng of 1-naphthaleneacetic acid (NAA) as an internal standard. The samples were centrifuged at 15,400 rpm for 5 minutes, and the supernatants were subjected to LC-MS/MS analysis equipped with the reverse-phase column (CORTECS UPLC Phenyl 1.6 μm, ϕ 2.1 × 75 mm; Waters). The peak area of (+)-GR24 and ABC-OH (degradation product) relative to that of NAA in extracted ion chromatogram were calculated. Detailed information about the analytical condition was described in table S2.

Fluorescence polarization

OPDA-PEG3-FL (100 nM) and 6×His-tagged OmKAI2d proteins were mixed with or without test chemicals on a half-area black 96-well plate (Greiner). The assays were performed in 20 μL of 1×PBS (−) containing 0.025% tween 20, 0.1% DMSO and 1% acetone. Measurements were taken on a microplate reader Infinite 200 PRO F Plex (TECAN) with the following settings (excitation 485(20)nm, emission 535(25)nm) at 25°C. The *G*-factors were determined using 100 nM OPDA-PEG3-FL in the assay buffer and adjusted so that the mP value of the free fluorescent probe as 20 units. kinetic measurements were performed at 5 min intervals.

Isothermal titration calorimetry

Isothermal titration calorimetry (ITC) experiments were performed in a MicroCal ITC200 MicroCalorimeter. Recombinant OmKAI2d3 (no tag) protein was prepared as previously described (*29*). The ligand at a concentration of 1 mM was titrated into a 40 μM protein solution in 20 steps of 2 μL and in 240-second intervals. Thermodynamic parameters were then calculated using the MicroCal ITC software as part of Origin (Originlab).

TR-FRET assay

TR-FRET assays were performed as previously described, with slight modifications (*33*). The experiments were conducted in HTRF 96-well low-volume white plate (Revvity) in 20 μL of reaction buffer [50 mM HEPES-KOH (pH7.0), 150 mM NaCl, 2 mM sodium ascorbate, 0.1 mg/mL BSA, 1× HTRF mAb anti-FLAG Tb-Conjugate (Revvity)] with OmKAI2d3/4/9/10-mEGFP (100 nM), 3×FLAG-OmSMAX1$_{D1M}$-6×His and indicated concentrations of chemicals from the acetone stock solution. The final concentration of acetone was adjusted to 2.1% (v/v). After the FLAG-tagged protein was incubated in reaction buffer at room temperature for 30 min, the SL receptor proteins and chemical solutions were added to a reaction mixture on ice. Measurements were taken on a microplate reader Infinite 200 PRO F Plex (TECAN) with the following settings (excitation 340(35)nm, emission A 520(10)nm, emission B 485(20)nm, 120 μs delay time, 200 μs integration time, 5 min interval). The TR-FRET signal was taken as the emission A/emission B intensity ratio.

Pull-down assay

Pull-down assays were performed according to the previous study, with slight modifications (*33*). OmKAI2d3/9-mEGFP (~6.3 μg) was immobilized on 5 or 10 μL of GFP-Trap Magnetic Particles M-270 (Proteintech) at 4℃ for 1 h with agitation. The beads were washed with 500 μL of washing buffer [50 mM HEPES-KOH (pH7.0), 150 mM NaCl, 0.1 mg/mL BSA, 10% glycerol, 0.1 % Tween 20] for 3 times. OmSMAX1$_{D1M}$ protein (0.01 μg) was added to the beads in 40 μL of assay buffer [50 mM HEPES-KOH (pH7.0), 150 mM NaCl, 2 mM sodium ascorbate, 0.1 mg/mL BSA, 10% glycerol, 1% DMSO, 0.1% Tween 20] with or without test chemicals. After incubation at 4℃ for 120 min with agitation, the beads were washed by 200 μL of washing buffer for 5 times at room temperature. Bound proteins were extracted by 20 μL of 1×SDS-PAGE sample buffer with boiling. The samples were subjected to SDS-PAGE and western blotting analysis.

OmKAI2d3/9-mEGFP (~6.3 μg) was immobilized on 10 μL of GFP-Trap Magnetic Particles M-270 at 4℃ for 1 h with agitation. The beads were washed with 500 μL of washing buffer [50 mM HEPES-KOH (pH7.0), 150 mM NaCl, 0.1 mg/mL BSA, 10% glycerol, 0.1 % Tween 20] for 3 times. OmMAX2 protein (24 μg) was added to the beads in 40 μL of assay buffer [50 mM HEPES-KOH (pH7.0), 150 mM NaCl, 2 mM sodium ascorbate, 0.1 mg/mL BSA, 10% glycerol, 1% DMSO, 0.1% Tween 20] with or without test chemicals. After incubation at room temperature for 180 min with agitation, the beads were washed by 200 μL of washing buffer for 5 times at room temperature. Bound proteins were extracted by 20 μL of 1×SDS-PAGE sample buffer with boiling. The samples were subjected to SDS-PAGE and western blotting analysis. GFP-tagged protein and FLAG-tagged protein were detected by western blotting according to the previous study (*33*).

Analysis of the covalent bound formation by non-reducing SDS-PAGE

6×His-OmKAI2d3 and 6×His-OmKAI2d3$^{S95A}$ (10 μM) were incubated with OPDA-PEG3-FL (10 μM) in the presence or absence of test chemicals in 20 μL of 1×PBS (−) containing 1% DMSO and 1% acetone at 25°C for 1 hour. The reaction was stopped by adding 20 μL of 2×SDS-PAGE sample buffer without reducing agents. The samples were subjected to 12% gel SDS-PAGE analysis. Images were obtained using the iBright CL 1500 (Invitrogen).

LC-MS/MS analysis of jasmonates from rice hydroponic culture media

We used *oscerk1* rice plants and the corresponding wild-type (*Oryza sativa* cv. Nipponbare BL no. 2) (*42*). Sterilized rice seeds were incubated at 30°C under dark conditions until radicle emergence. The seeds were transferred onto agar media (*58*) and grown under long day conditions (16 h light at 28 °C/8 h dark at 25°C) for 5-6 days. Rice plants were additionally cultivated in hydroponic culture system under sufficient phosphate conditions (600 μM Pi) for 7 days (*59*). Hydroponic culture media was renewed on a day before elicitor treatment. 250 μL of chitin oligosaccharides solution (100 mg/mL) or distilled water (as control) was supplemented into rice hydroponic culture (approximately 10 mL). After the incubation at room temperature for 2 hours, hydroponic culture media was collected from one or two tubes and spiked with internal standards, OPDA *d*6, JA *d*6, JA-Ile *d*6. Root exudates were extracted with ethyl acetate twice. The organic phase was concentrated under a stream of nitrogen. The residue was dissolved in acetonitrile and subjected to LC-MS/MS analysis equipped with the reverse-phase column (CORTECS UPLC Phenyl 1.6 μm, ϕ 2.1 × 75 mm; Waters). Detailed information about the analytical condition was described in table S3.

Germination assay using extracts of rice hydroponic culture media

We used *d10-1*and *cpm2* rice plants, as well as the corresponding wild-type of *cpm2* (*Oryza sativa* cv. Nihonmasari) (*45*, *60*). Rice plants were grown and treated with chitin oligosaccharides as described above. Hydroponic culture media were collected from two tubes and extracted with ethyl acetate twice. The organic phase was concentrated under a stream of nitrogen. The residue was dissolved in sterilized water and used for the germination assay. At the same time, quantification of jasmonates in hydroponic culture media was conducted with the same procedure as described above. Because the amount of *cis*-OPDA was below the detection limit of LC-MS/MS analysis, it was not measured in the experiments which were conducted in parallel with the germination assay.
Note: Because the homozygous *cpm2* mutant plants are male sterile, seeds were obtained from the heterozygous plants. PCR-based genotyping was performed using primers (cpm2-F: 5′-CGCTCGTTGATCTCGTACAC-3′ and cpm2-R: 5′-GTCGCTGTTCTCGCCGAAG-3′) and the homozygous *cpm2* mutant plants were identified and used for the experiments.

Data analysis and graphics

For the calculations of $EC_{50}$ values, the plots were fitted by a three or four-parameter logistic model using the R drc package (*61*). Graphs were drawn using the R ggplot2 package (*62*).

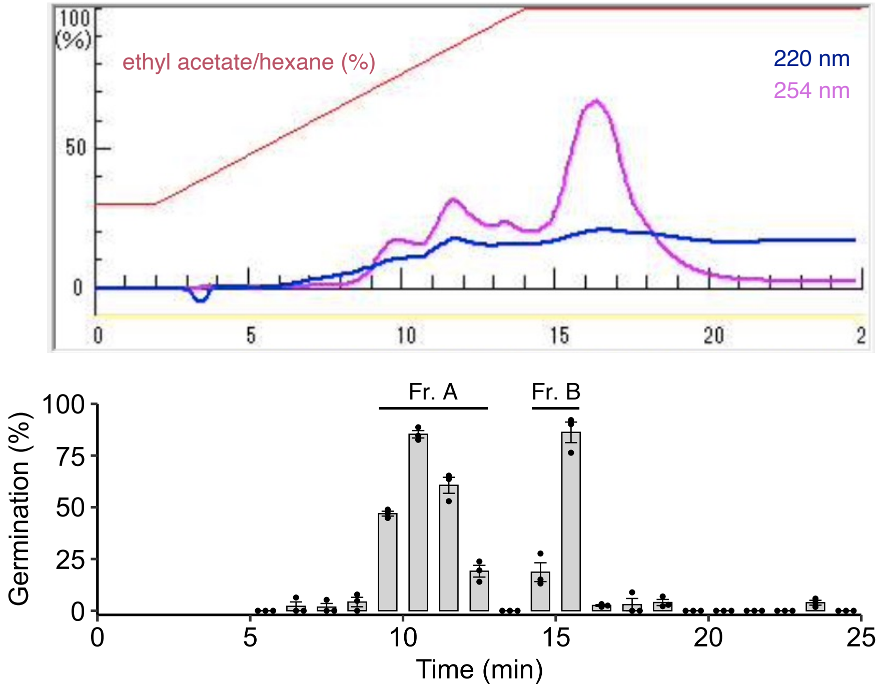


**Fig. S1. Purification of the active compounds from ethyl acetate extracts.** Chromatogram of the ethyl acetate extracts from *G. fujikuroi* culture filtrates (upper panel). Germination induing activity of the collected fractions was evaluated by *O. minor* seed germination assay. The assay was performed with each fraction concentrated 100-fold relative to the original culture volume. Data are presented as the means ± SE of three biological replicates. Dots represent individual data points.

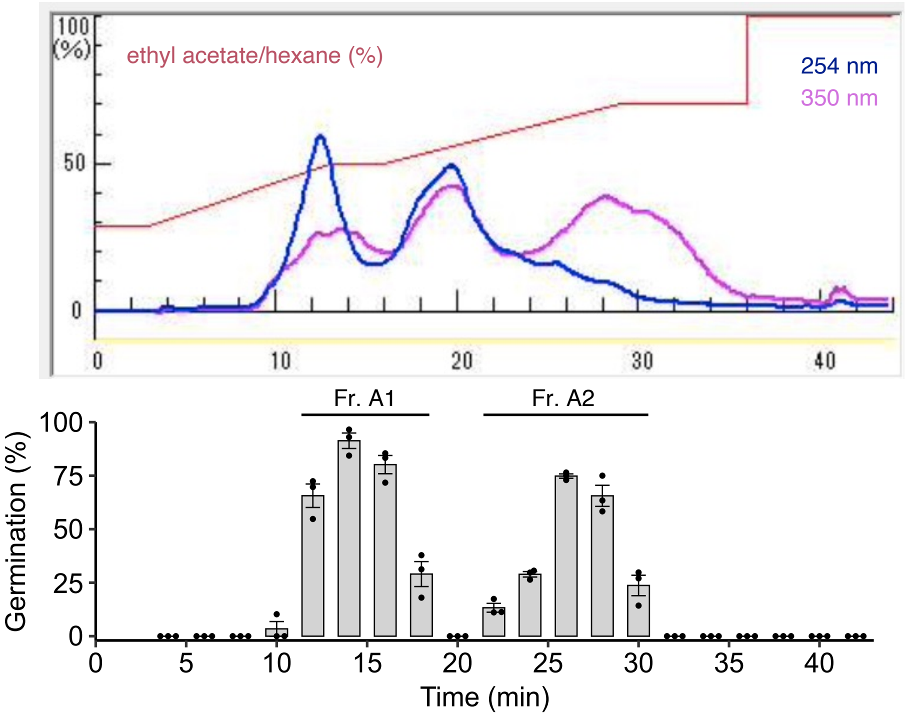


**Fig. S2. Purification of the active compounds from the fraction Fr. A.** Chromatogram of Fr. A (upper panel). Germination induing activity of the collected fractions was evaluated by *O. minor* seed germination assay. The assay was performed with each fraction concentrated 100-fold relative to the original culture volume. Data are presented as the means ± SE of three biological replicates. Dots represent individual data points.

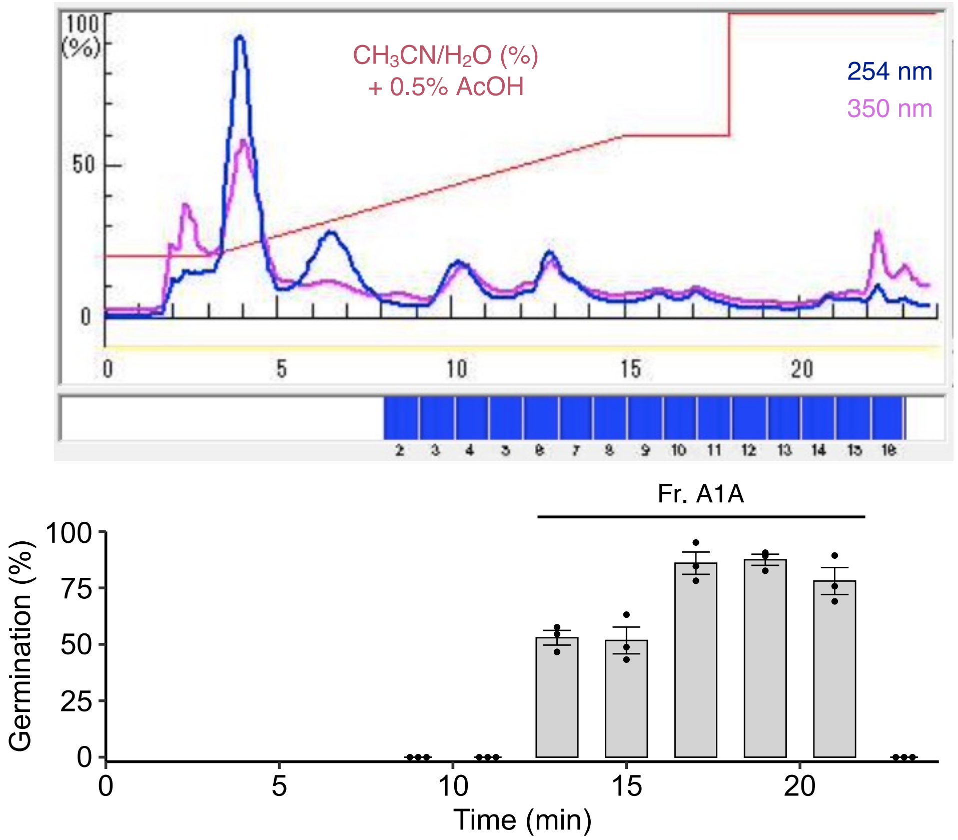


**Fig. S3. Purification of the active compounds from the fraction Fr. A1.** Chromatogram of Fr. A1 (upper panel). Germination induing activity of the collected fractions was evaluated by *O. minor* seed germination assay. The assay was performed with each fraction concentrated 1000-fold relative to the original culture volume. Data are presented as the means ± SE of three biological replicates. Dots represent individual data points.

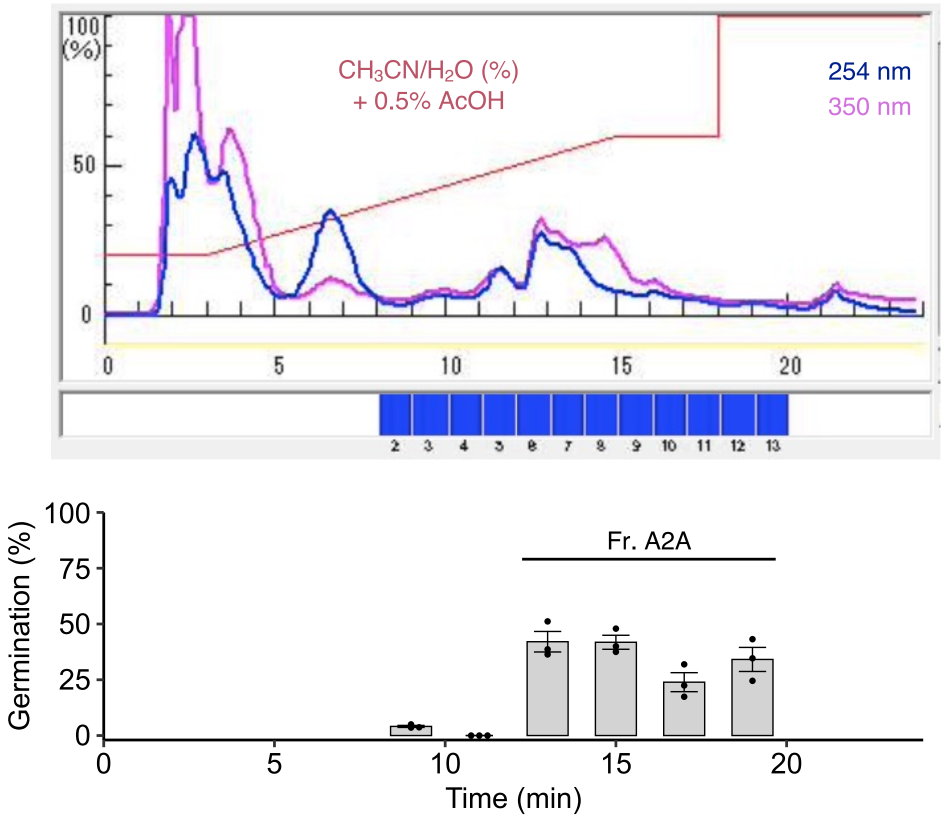


**Fig. S4. Purification of the active compounds from the fraction Fr. A2.** Chromatogram of Fr. A2 (upper panel). Germination induing activity of the collected fractions was evaluated by *O. minor* seed germination assay. The assay was performed with each fraction concentrated 1000-fold relative to the original culture volume. Data are presented as the means ± SE of three biological replicates. Dots represent individual data points.

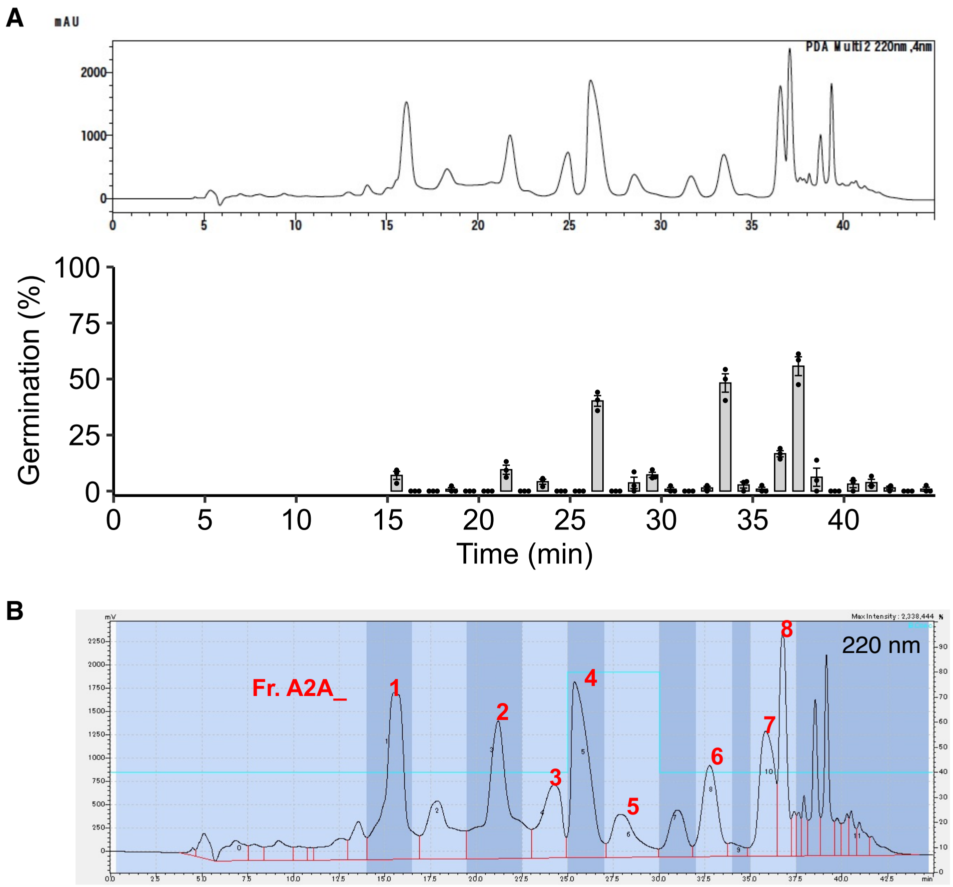


**Fig. S5. Purification of the active compounds from the fraction Fr. A2A.** (**A**) Chromatogram of Fr. A2A (upper panel). Germination induing activity of the collected fractions was evaluated by *O. minor* seed germination assay. The assay was performed with each fraction concentrated 1000-fold relative to the original culture volume. Data are presented as the means ± SE of three biological replicates. Dots represent individual data points. (**B**) Fractions were pooled as indicated by different colors.

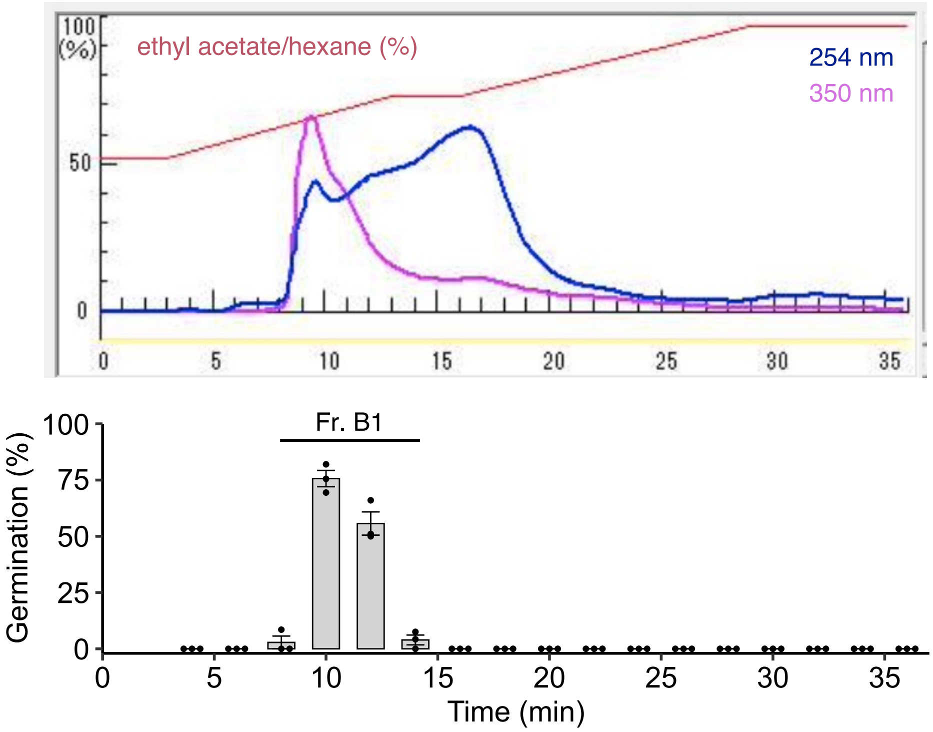


**Fig. S6. Purification of the active compounds from Fr. B.** Chromatogram of Fr. B (upper panel). Germination induing activity of the collected fractions was evaluated by *O. minor* seed germination assay. The assay was performed with each fraction concentrated 100-fold relative to the original culture volume. Data are presented as the means ± SE of three biological replicates. Dots represent individual data points.

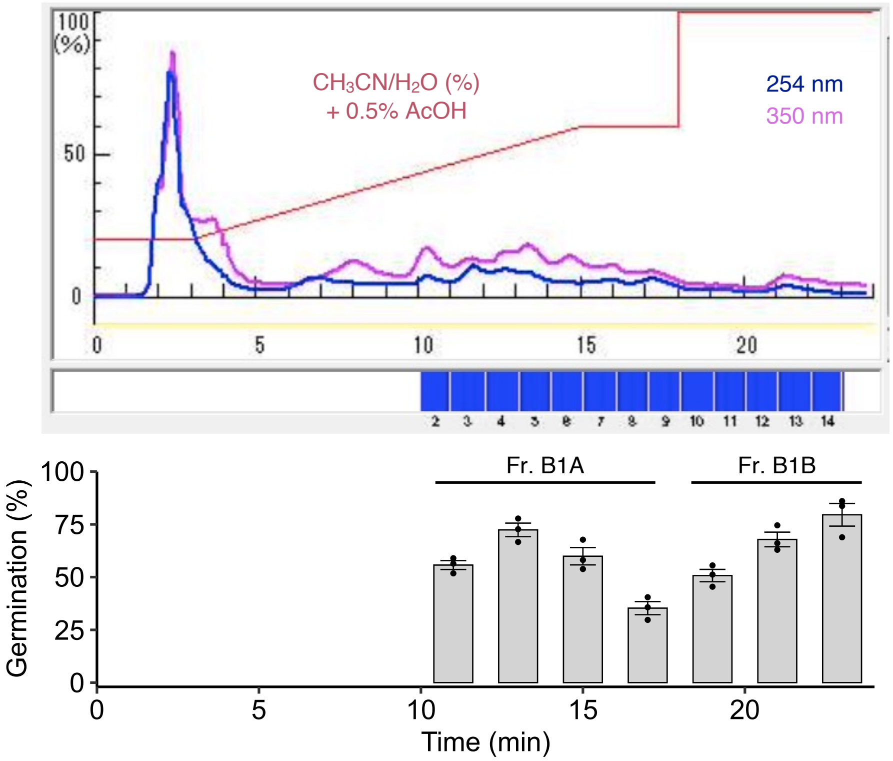


**Fig. S7. Purification of the active compounds from Fr. B1.** Chromatogram of Fr. B1 (upper panel). Germination induing activity of the collected fractions was evaluated by *O. minor* seed germination assay. The assay was performed with each fraction concentrated 100-fold relative to the original culture volume. Data are presented as the means ± SE of three biological replicates. Dots represent individual data points.

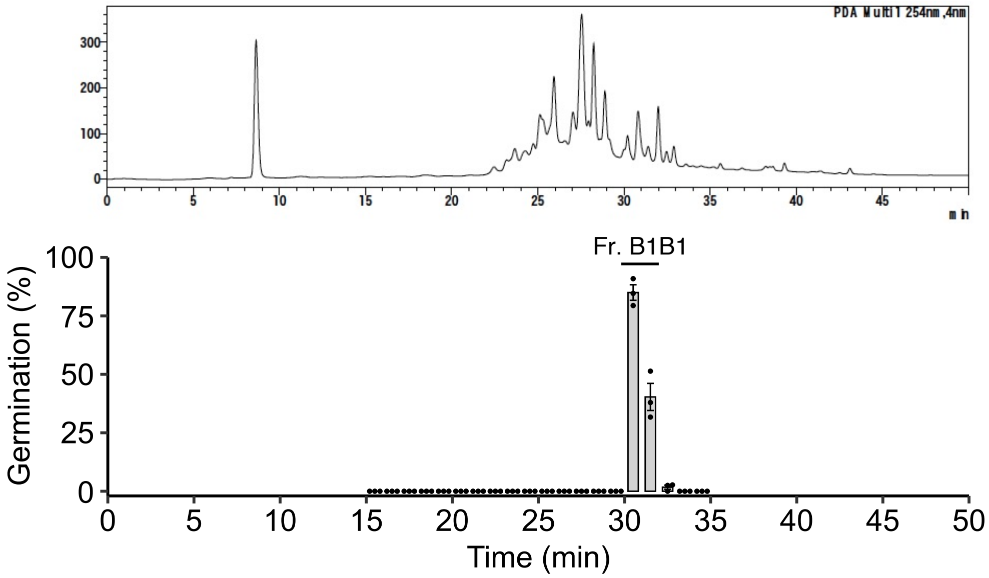


**Fig. S8. Purification of the active compounds from Fr. B1B.** Chromatogram of Fr. B1B (upper panel). Germination induing activity of the collected fractions was evaluated by *O. minor* seed germination assay. The assay was performed with each fraction concentrated 1000-fold relative to the original culture volume. Data are presented as the means ± SE of three biological replicates. Dots represent individual data points.

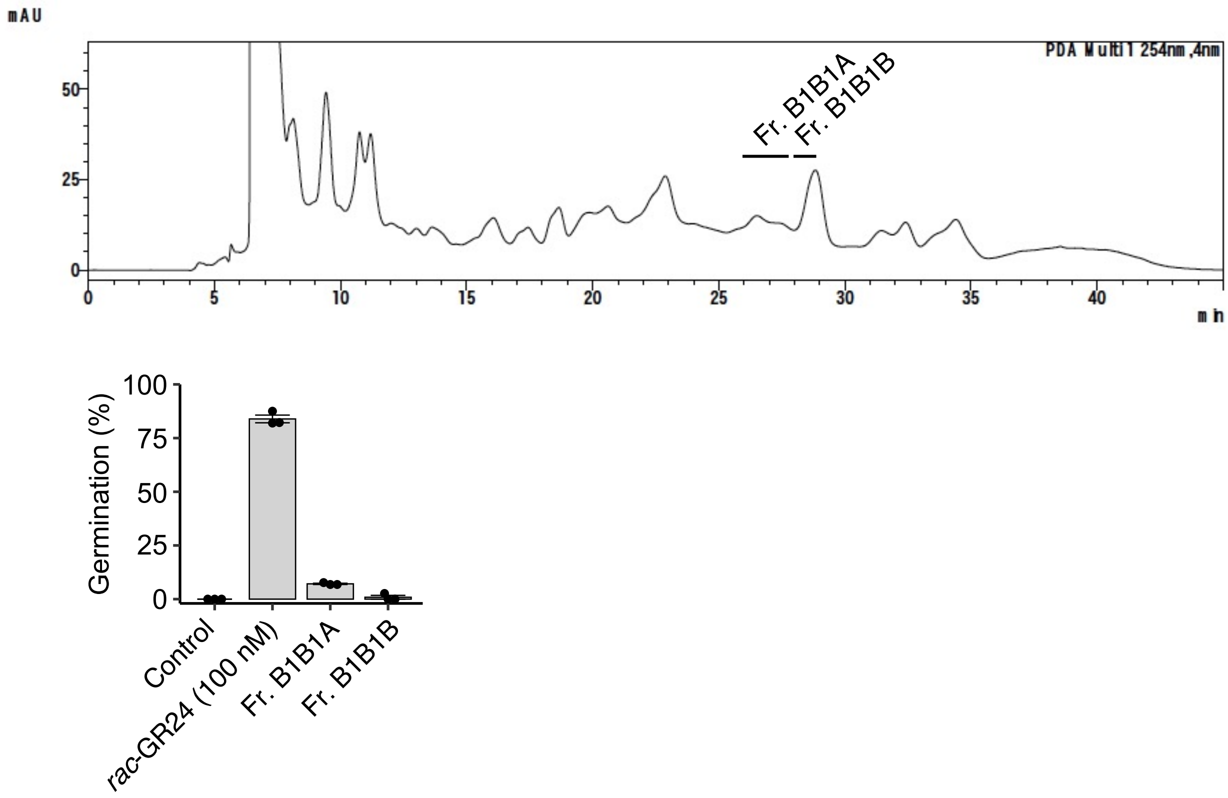


**Fig. S9. Purification of the active compounds from Fr. B1B1.** Chromatogram of Fr. B1B1 (upper panel). Germination induing activity of the collected fractions was evaluated by *O. minor* seed germination assay. The assay was performed with each fraction concentrated 100-fold relative to the original culture volume. Data are presented as the means ± SE of three biological replicates. Dots represent individual data points.

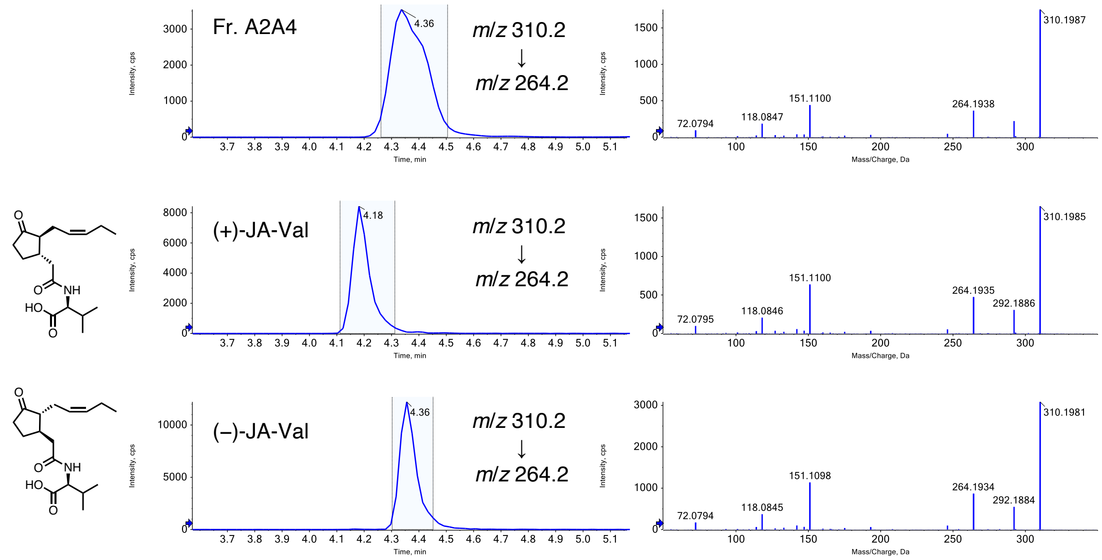


**Fig. S10. LC-MS/MS analysis of Fr. A2A4.** LC-MS/MS chromatogram of Fr. A2A4 and chemically synthesized JA-Val isomers in the multiple reaction monitoring mode. Selected reaction monitoring (*m*/*z* 310.2 > *m*/*z* 264.2) (left panel). Full-scan spectra of fragment ions derived from the precursor ion (*m*/*z* 310.2) of the left panel peak highlighted in pale blue (right panel).

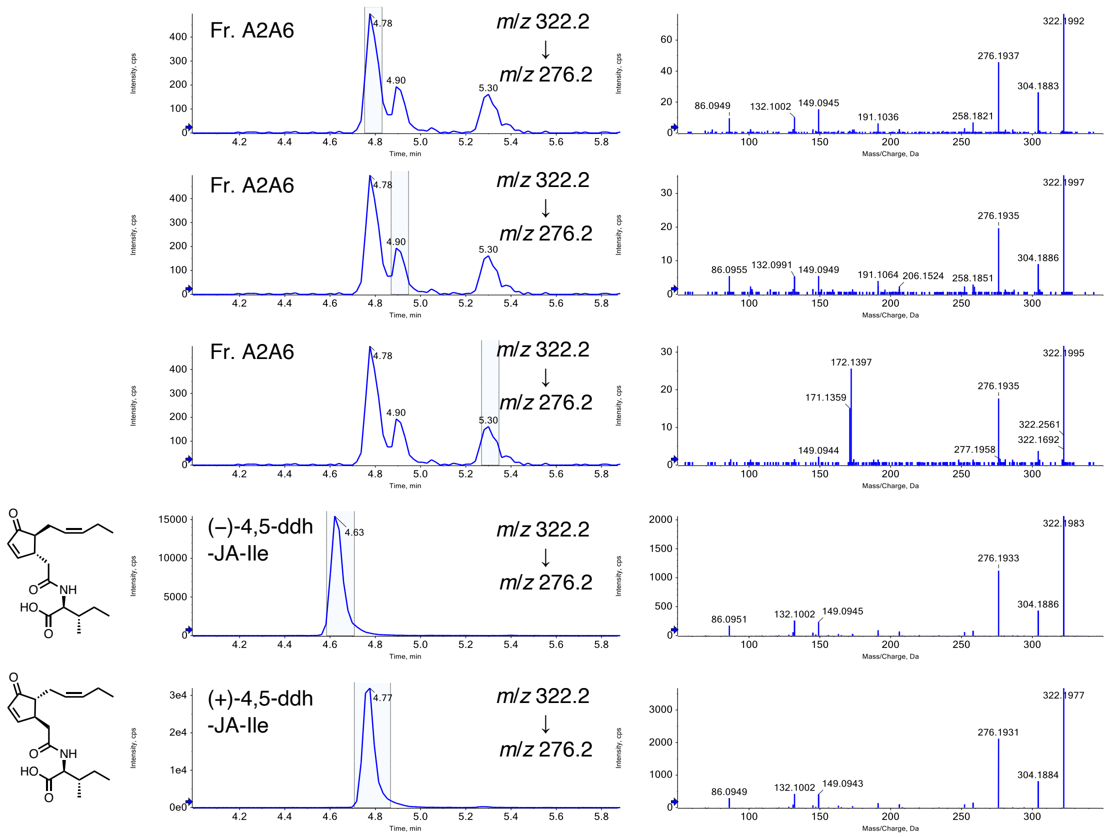


**Fig. S11. LC-MS/MS analysis of Fr. A2A6.** LC-MS/MS chromatogram of Fr. A2A6 and chemically synthesized ddh-JA-Ile isomers in the multiple reaction monitoring mode. Selected reaction monitoring (*m*/*z* 322.2 > *m*/*z* 276.2) (left panel). Full-scan spectra of fragment ions derived from the precursor ion (*m*/*z* 322.2) of the left panel peak highlighted in pale blue (right panel).

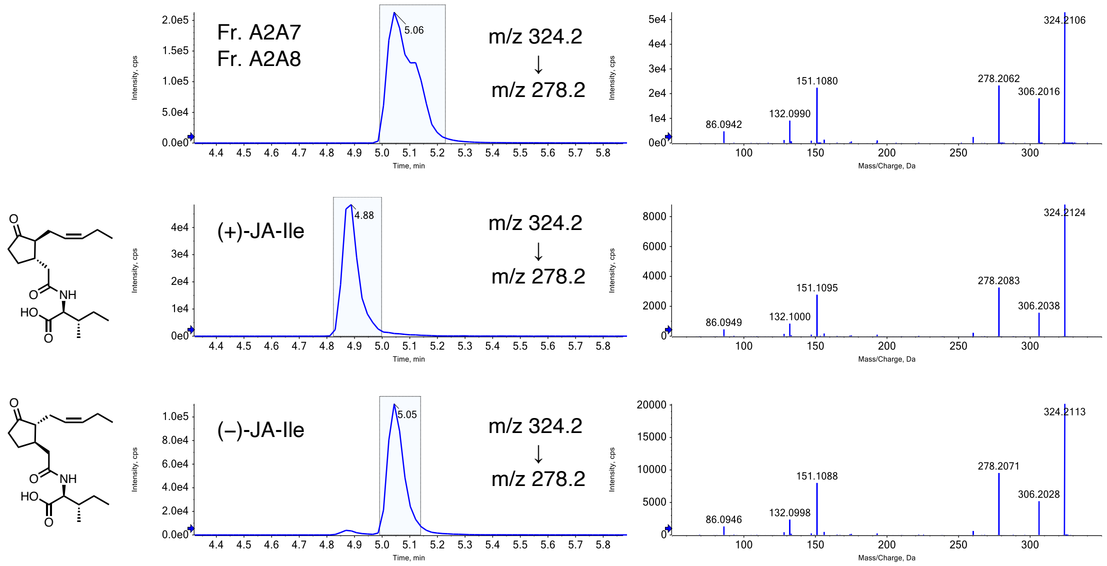


**Fig. S12. LC-MS/MS analysis of Fr. A2A7/A2A8.** LC-MS/MS chromatogram of Fr. A2A7/A2A8 and chemically synthesized JA-Ile isomers in the multiple reaction monitoring mode. Selected reaction monitoring (JA-Ile: *m*/*z* 324.2 > *m*/*z* 278.2) (left panel). Full-scan spectra of fragment ions derived from the precursor ion (JA-Ile: *m*/*z* 322.2) of the left panel peak highlighted in pale blue (right panel).

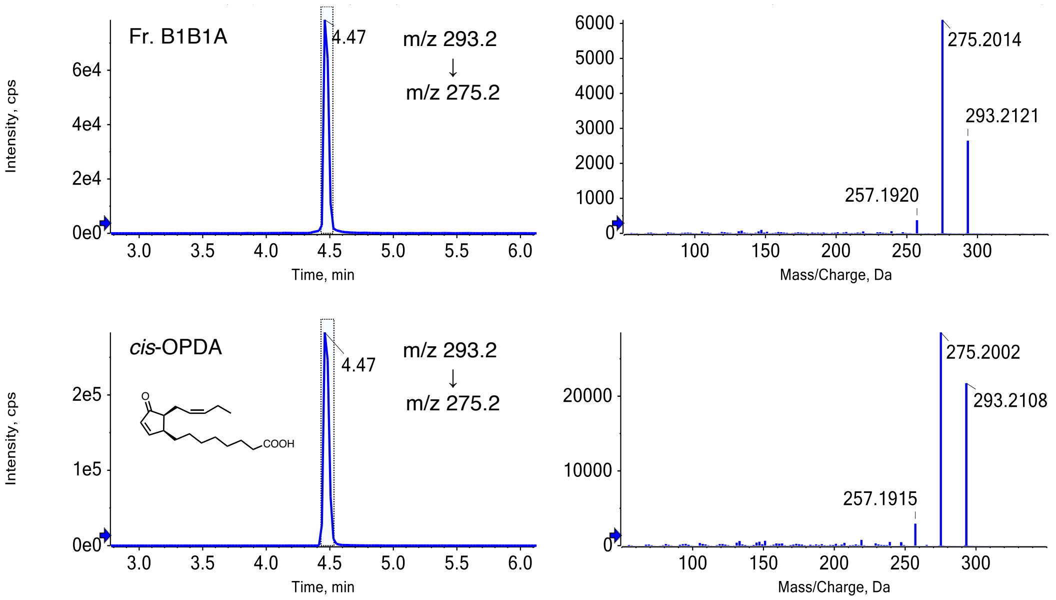


**Fig. S13. LC-MS/MS analysis of Fr. B1B1A.** LC-MS/MS chromatogram of Fr. B1B1A and chemically synthesized *cis*-OPDA in the multiple reaction monitoring mode. Selected reaction monitoring (*m*/*z* 293.2 > *m*/*z* 275.2) (left panel). Full-scan spectra of fragment ions derived from the precursor ion (*m*/*z* 310.2) of the left panel peak highlighted in pale blue (right panel).

**A**

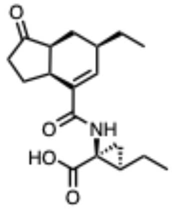

**B**

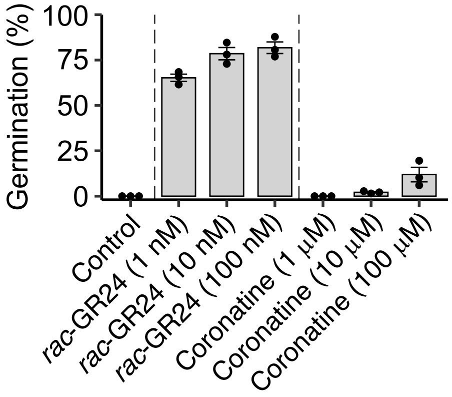


**Fig. S14. Germination inducing activity of coronatine toward *O. minor* seeds.** (**A**) Chemical structure of coronatine. (**B**) Germination rate of *O. minor* seeds after treatment with test chemicals. Data are presented as the means ± SE of three biological replicates. Dots represent individual data points.

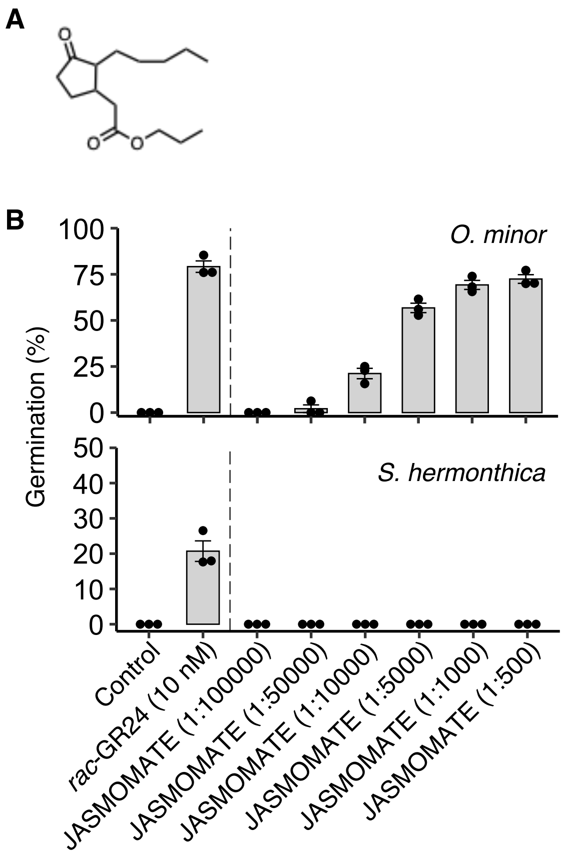


**Fig. S15. Germination inducing activity of JASMOMATE toward *O. minor* and *S. hermonthica* seeds.** (**A**) Chemical structure of propyl dihydrojasmonate. (**B**) Germination rate of *O. minor* and *S. hermonthica* seeds after treatment with test chemicals. Data are presented as the means ± SE of three biological replicates. Dots represent individual data points.

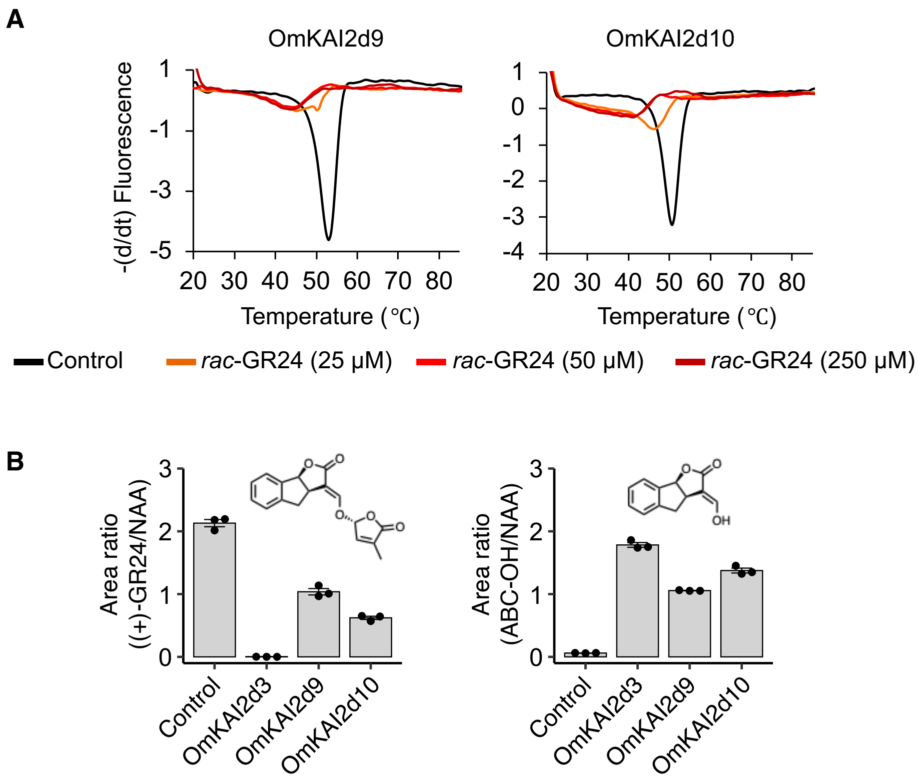


**Fig. S16. Evaluation of the SL binding and hydrolysis ability of OmKAI2d9/10.** (A) Melting temperature of OmKAI2d9/10 in the presence or absence of *rac*-GR24. Data are presented as the means of three technical replicates. (B) OmKAI2d-mediated hydrolysis of (+)-GR24. Relative amount of the remaining substrate (left panel) and the hydrolysis product (right panel) were measured by LC-MS/MS. Control was performed without proteins. Data are presented as the means ± SE of three technical replicates. Dots represent individual data points.

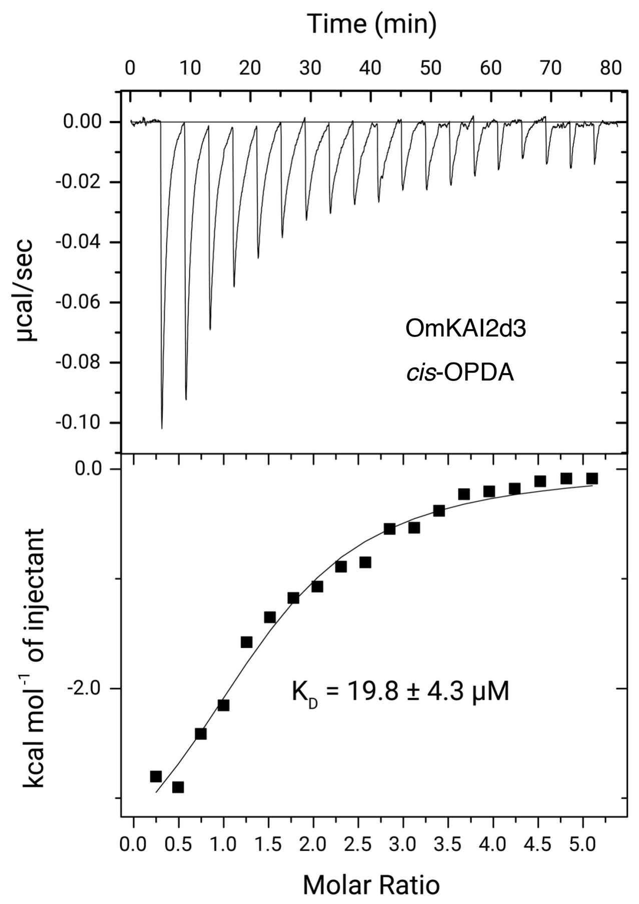


**Fig. S17. Dissociation constant of *cis*-OPDA to OmKAI2d3 measured by ITC.** *cis*-OPDA (mM) was injected into OmKAI2d3 solution in the ITC cell.

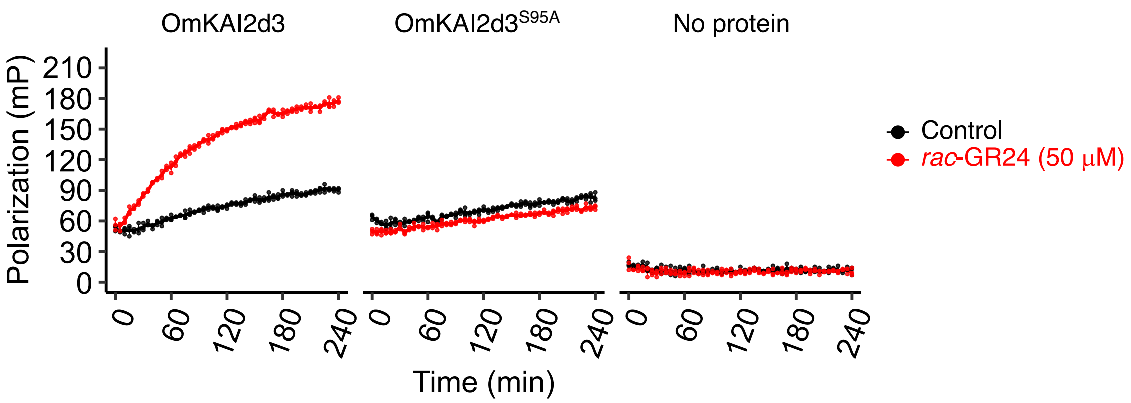


**Fig. S18. Kinetic FP assay using OmKAI2d3$^{S95A}$.** The assays were performed using OPDA-PEG3-FL (100 nM) and OmKAI2d3/OmKAI2d3$^{S95A}$ (10 mM). Control was performed without *rac*-GR24. Data are presented as the means ± SE of three technical replicates. Dots represent individual data points.

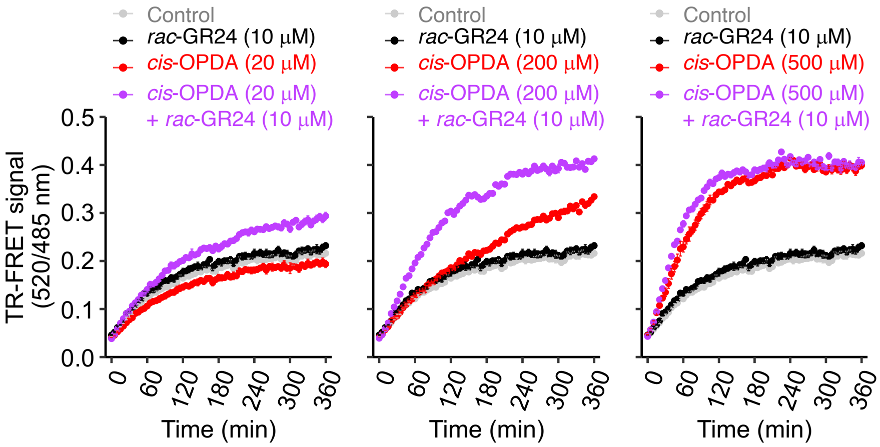


**Fig. S19. Kinetic measurements of OmKAI2d9–OmSMAX1$_{D1M}$ TR-FRET signals in the presence of *cis*-OPDA and *rac*-GR24.** Control was performed without test chemicals. Data are presented as the means ± SE of three technical replicates. Data at 300 min are provided in Fig. 4D for comparison.

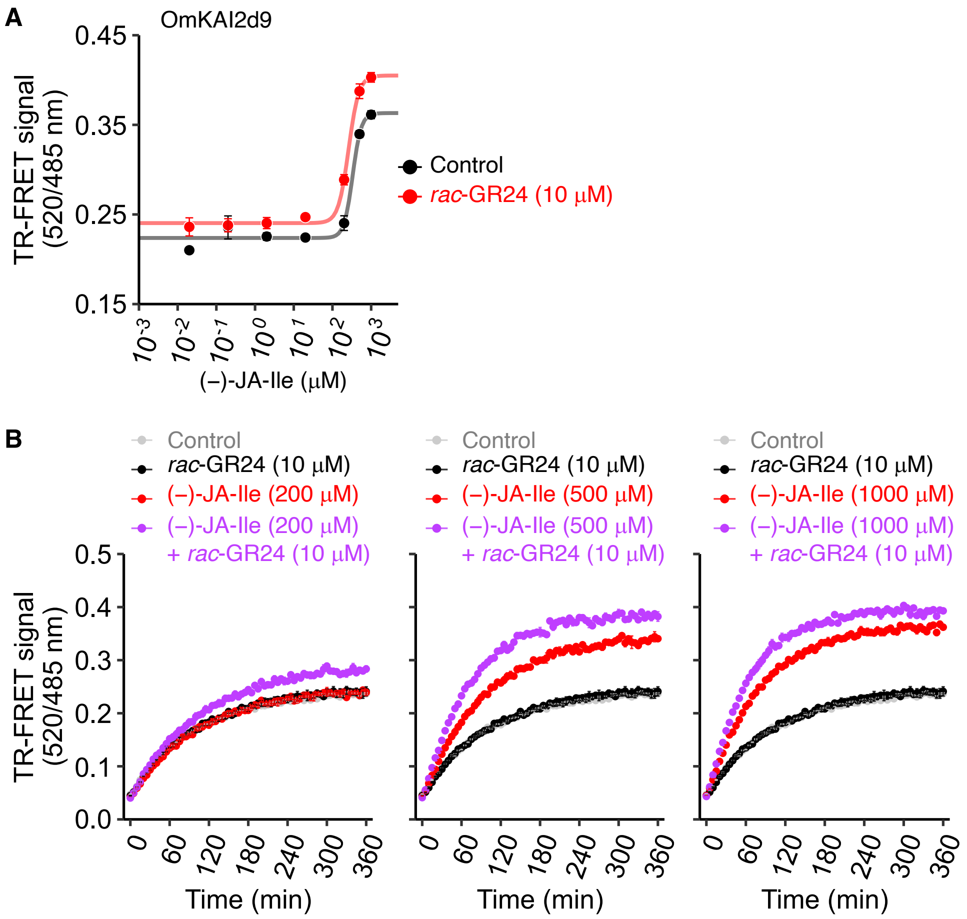


**Fig. S20. TR-FRET assay using OmKAI2d9–OmSMAX1$_{D1M}$ in the presence of (−)-JA-Ile and *rac*-GR24.** (A) Dose-titration of (−)-JA-Ile in the presence or absence of *rac*-GR24. (B) Kinetic measurements of TR-FRET signals. Control was performed without test chemicals. Data are presented as the means ± SE of three technical replicates.

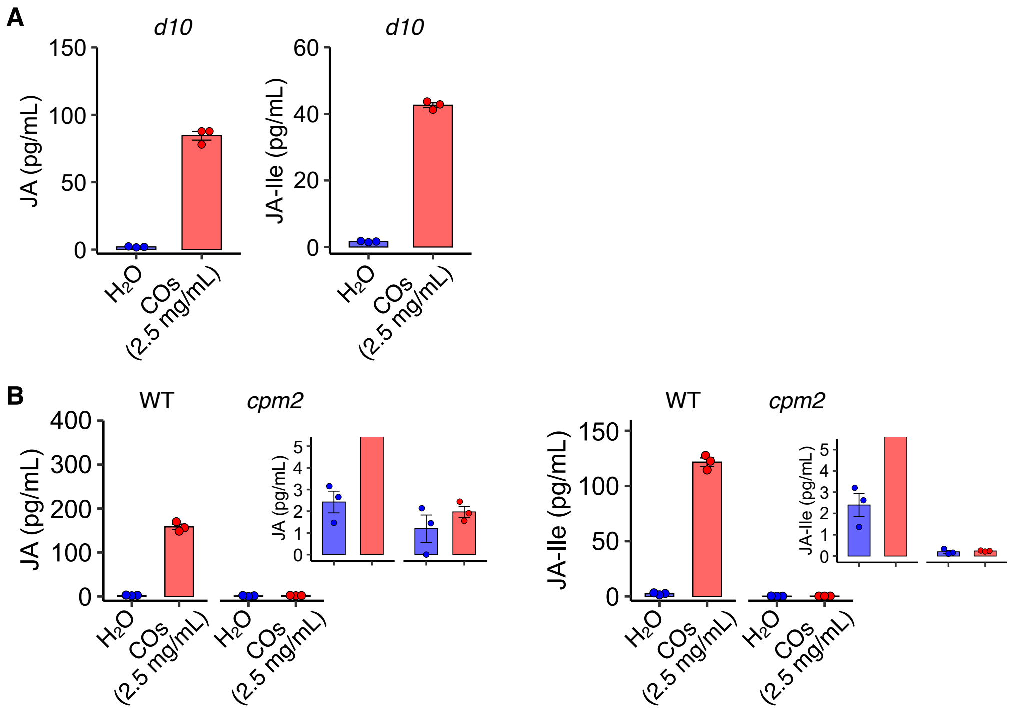


**Fig. S21. Effect of COs on jasmonate-exudation in rice hydroponic culture of the SL-biosynthesis and jasmonate-biosynthesis mutants.** LC-MS/MS analysis of JA and JA-Ile in hydroponic cultures of (A) *d10* or (B) *cpm2* and corresponding wild-type plants after treatment with COs elicitor for 2 hours. The inset (upper right) shows an enlarged view of the graph. Data are presented as the means ± SE of three biological replicates. Dots represent individual data points.

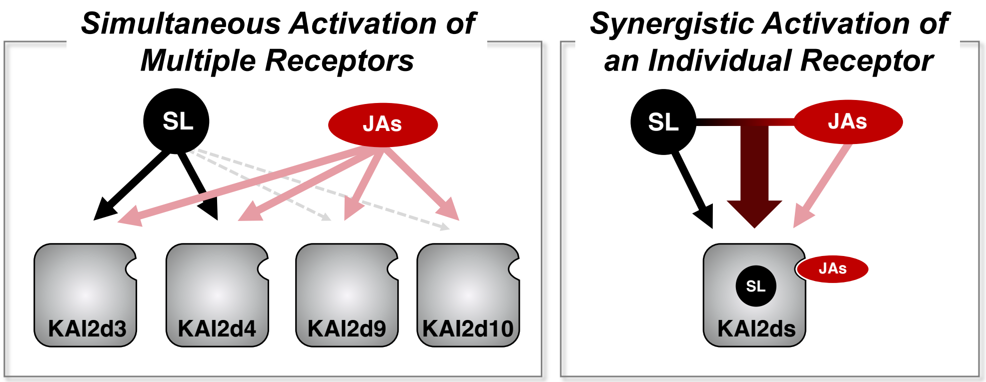


**Fig. S22. Putative models of the synergistic effect of SLs and jasmonates.** SLs strongly activate several KAI2d proteins. Jasmonates (JAs) weakly activate broad range of KAI2d proteins. The combination of SLs and jasmonates results in a potent and broad-spectrum activation of duplicated KAI2d proteins (left panel). Jasmonates allosterically interact with KAI2d proteins and synergistically activate the receptor with SLs (right panel).

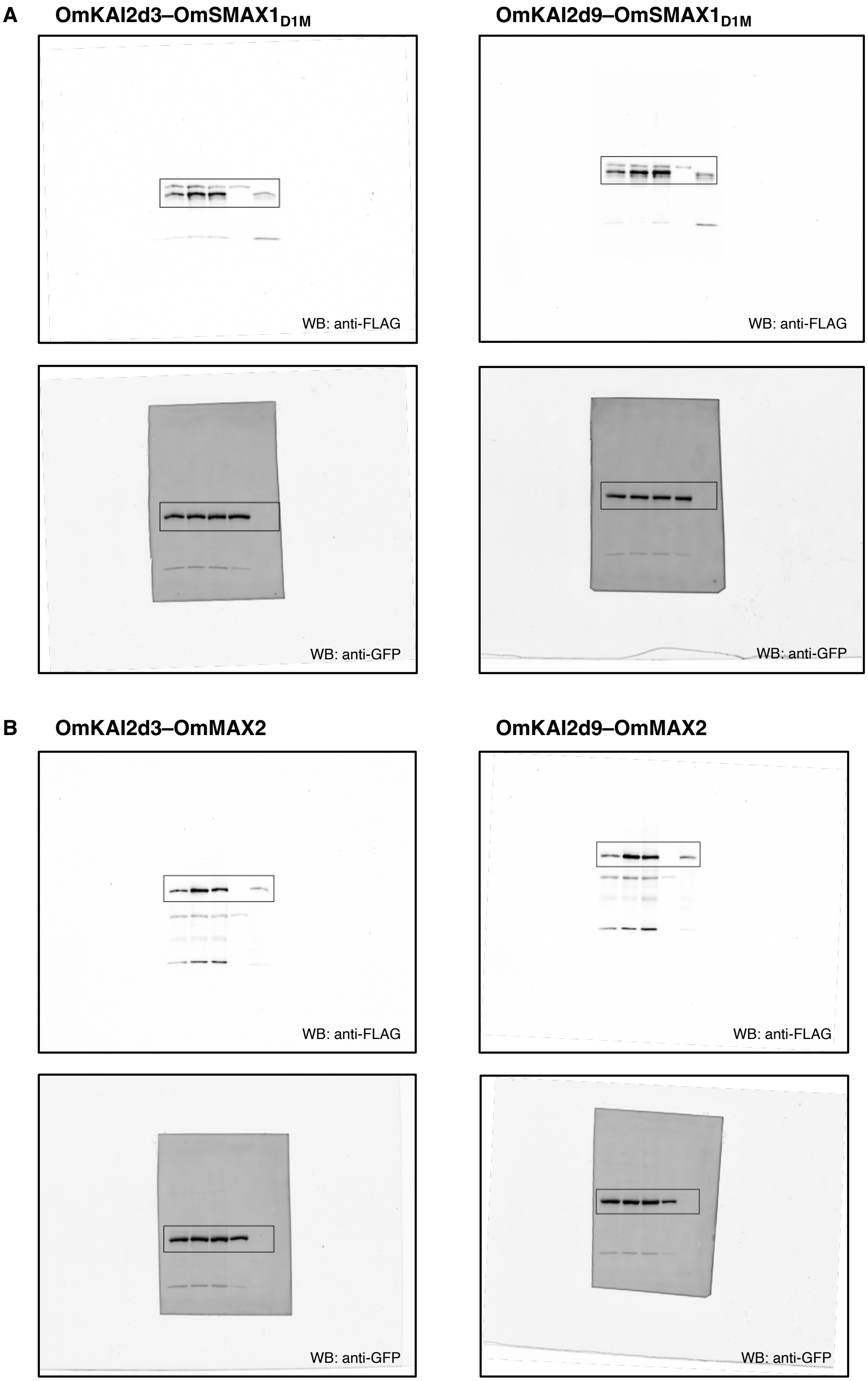


**Fig. S23. Uncropped images of the immunoblots shown in (A) Fig. 3F and (B) Fig. 3G.**

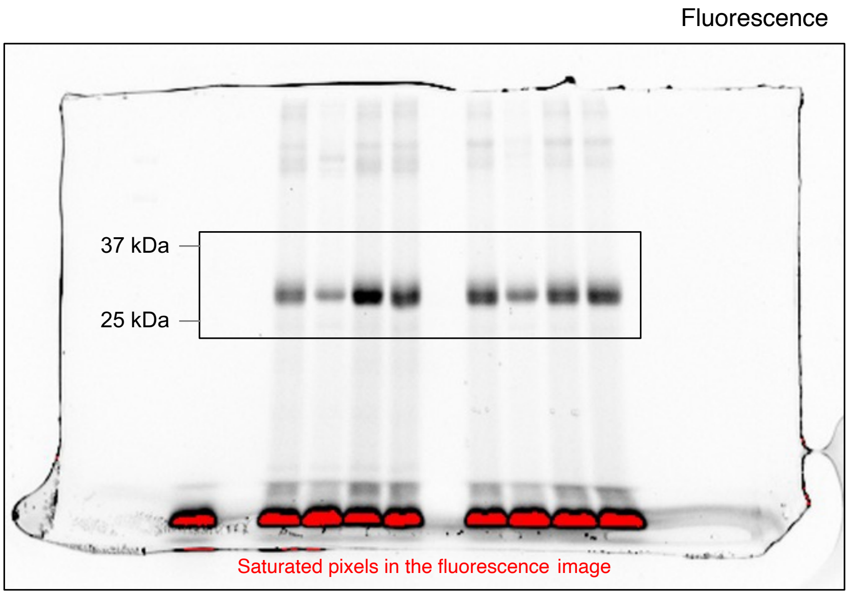


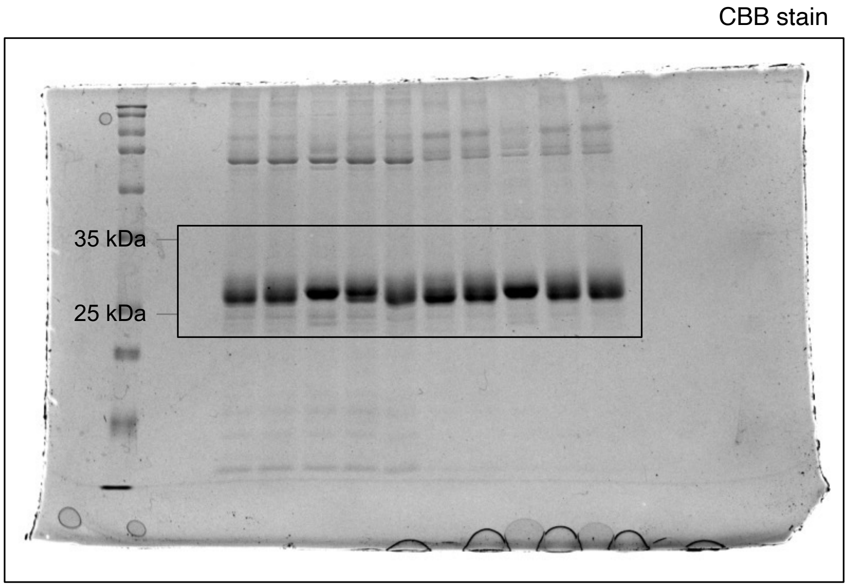


**Fig. S24. Uncropped images of the gels shown in Fig. 4B.**

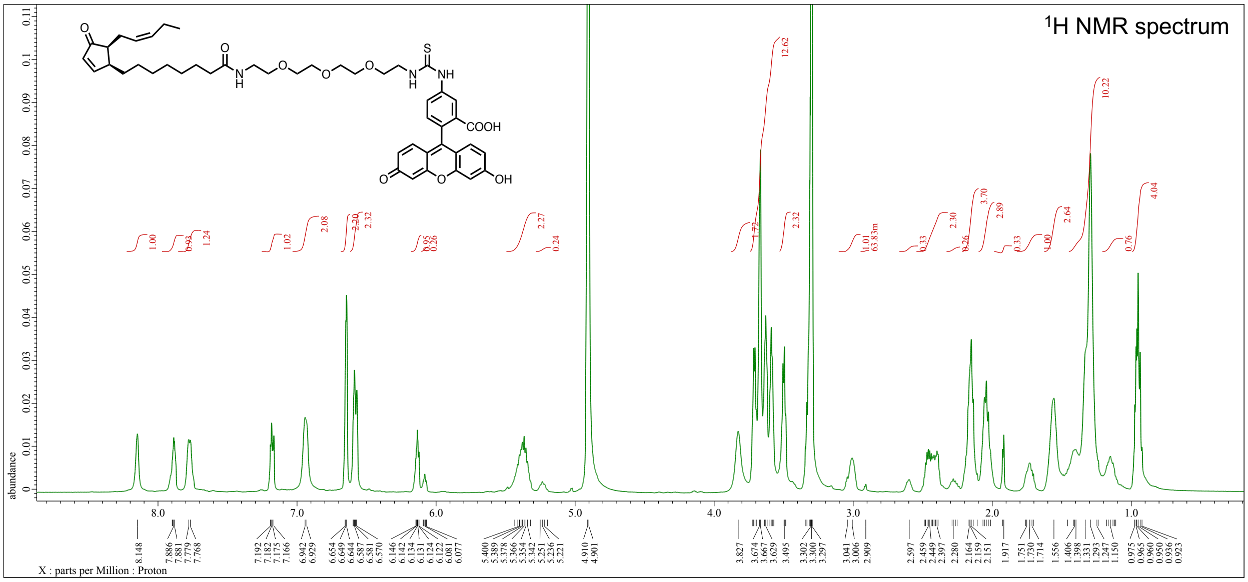


**Fig. S25. [1]H NMR spectrum of OPDA-PEG3-FL ($CD_3OD$, 500 MHz).**

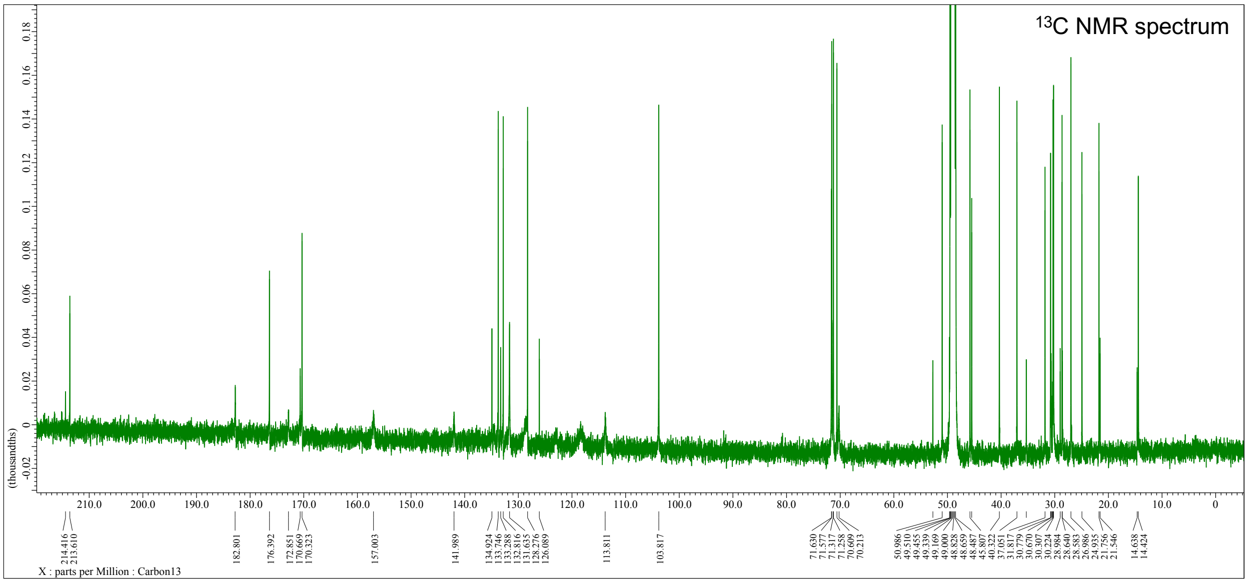


**Fig. S26.** $^{13}C$ **NMR spectrum of OPDA-PEG3-FL (**$CD_3OD$**, 125 MHz).**

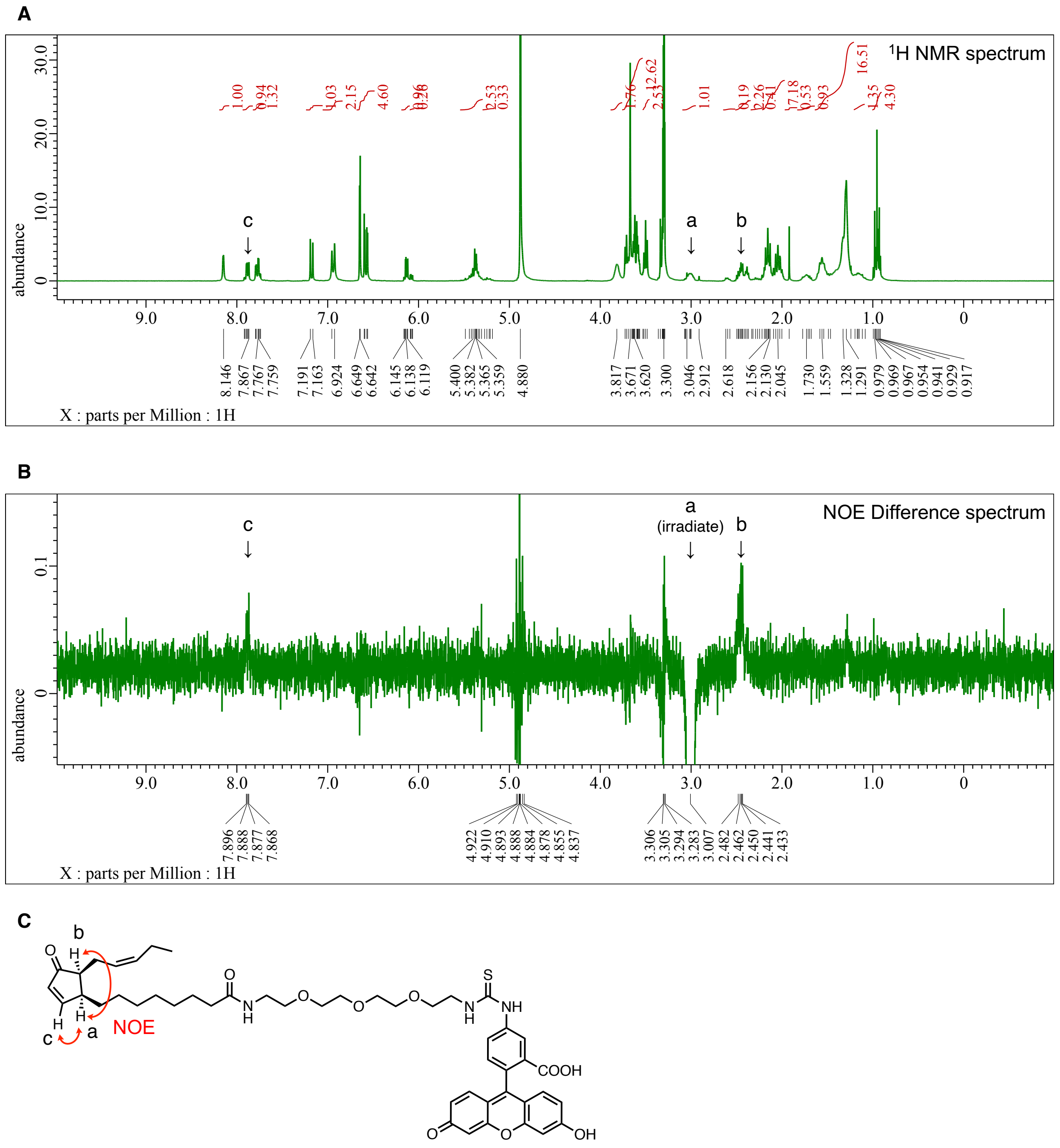


**Fig. S27. Differential NOE experiment on OPDA-PEG3-FL ($CD_3OD$, 300 MHz).** (A) $^1$H MNR spectrum of OPDA-PEG3-FL. (B) NOE Difference spectrum of OPDA-PEG3-FL. The proton signal at around 3 ppm was selectively irradiated. (C) Observed NOE correlations between the indicated protons in OPDA-PEG3-FL.

**Table S1. LC-MS/MS analytical conditions for the structural identification of the active compounds.**

MS conditions

| Compound name | MW | Parent ion (*m*/*z*) | Fragment ion (*m*/*z*) used for peak extraction | Declustering potential (V) | Collision energy (V) |
|---|---|---|---|---|---|
| (+/−)-JA-Val | 309 | 310 | 264.19 | 80 | 15 |
| (+/−)-JA-Ile | 323 | 324 | 278.20 | 80 | 15 |
| (+/−)-ddh-JA-Ile | 321 | 322 | 276.19 | 80 | 15 |
| (+)-*cis*-OPDA | 292 | 293 | 275.20 | 80 | 15 |

LC conditions (except for *cis*-OPDA)

| Time (min) | Flow rate (mL/min) | Solvent A (%) | Solvent B (%) |
|---|---|---|---|
| 0 | 0.3 | 80 | 20 |
| 5.00 | 0.3 | 50 | 50 |
| 5.01 | 0.3 | 2 | 98 |
| 7.00 | 0.3 | 2 | 98 |
| 7.01 | 0.3 | 80 | 20 |
| 13.00 | 0.3 | 80 | 20 |

Column: CORTECS UPLC Phenyl 1.6 mm, ϕ 2.1 × 75 mm (Waters)
Solvent A : Water (0.05% AcOH)
Solvent B : Acetonitrile (0.05% AcOH)

LC conditions (for *cis*-OPDA)

| Time (min) | Flow rate (mL/min) | Solvent A (%) | Solvent B (%) |
|---|---|---|---|
| 0 | 0.3 | 98 | 2 |
| 4.50 | 0.3 | 2 | 98 |
| 7.00 | 0.3 | 2 | 98 |
| 7.01 | 0.3 | 98 | 2 |
| 15.00 | 0.3 | 98 | 2 |

Column: CORTECS UPLC Phenyl 1.6 mm, ϕ 2.1 × 75 mm (Waters)
Solvent A : Water (0.05% AcOH)
Solvent B : Acetonitrile (0.05% AcOH)

### Table S2. LC-MS/MS analytical conditions for the GR24 hydrolysis assay.

MS conditions

| Compound name | MW | Parent ion (*m*/*z*) | Fragment ion (*m*/*z*) used for peak area calculation | Declustering potential (V) | Collision energy (V) |
|---|---|---|---|---|---|
| (+)-GR24 | 298 | 299 | 185.06 | 80 | 15 |
| ABC-OH | 202 | 203 | 185.06 | 80 | 15 |
| NAA | 186 | 187 | 141.07 | 80 | 15 |

LC conditions

| Time (min) | Flow rate (mL/min) | Solvent A (%) | Solvent B (%) |
|---|---|---|---|
| 0 | 0.3 | 80 | 20 |
| 5.00 | 0.3 | 2 | 98 |
| 7.00 | 0.3 | 2 | 98 |
| 7.01 | 0.3 | 80 | 20 |
| 13.00 | 0.3 | 80 | 20 |

Column: CORTECS UPLC Phenyl 1.6 mm, $\phi$ 2.1 × 75 mm (Waters)
Solvent A : Water (0.05% AcOH)
Solvent B : Acetonitrile (0.05% AcOH)

### Table S3. LC-MS/MS analytical conditions for the quantification of jasmonates.

MS conditions

| Compound name | MW | Parent ion (*m*/*z*) | Fragment ion (*m*/*z*) used for peak area calculation | Declustering potential (V) | Collision energy (V) |
|---|---|---|---|---|---|
| *cis*-OPDA | 292 | 291 | 165.13 | −80 | −35 |
| *trans*-OPDA *d*6 | 298 | 297 | 171.16 | −80 | −35 |
| JA | 210 | 209 | 59.01 | −80 | −35 |
| JA *d*6 | 216 | 215 | 59.01 | −80 | −35 |
| JA-Ile | 323 | 322 | 130.08 | −80 | −35 |
| JA-Ile *d*6 | 329 | 328 | 130.08 | −80 | −35 |

LC conditions

| Time (min) | Flow rate (mL/min) | Solvent A (%) | Solvent B (%) |
|---|---|---|---|
| 0 | 0.3 | 98 | 2 |
| 4.50 | 0.3 | 2 | 98 |
| 7.00 | 0.3 | 2 | 98 |
| 7.01 | 0.3 | 98 | 2 |
| 15.00 | 0.3 | 98 | 2 |

Column: CORTECS UPLC Phenyl 1.6 mm, ϕ 2.1 × 75 mm (Waters)
Solvent A : Water (0.05% AcOH)
Solvent B : Acetonitrile (0.05% AcOH)
